# A Scalable Design for Proximity-Inducing Molecules

**DOI:** 10.64898/2026.02.20.706349

**Authors:** Endri Karaj, Varsha Venkatarangan, Shaimaa H. Sindi, Surached Siriwongsup, Chaiheon Lee, Rajaiah Pergu, Vedagopuram Sreekanth, Karishma Kailass, Kien Tran, Prashant Singh, Sameek Singh, Junya Kawai, Jeffrey E. Fung, Mahilet Tefera, Rohil Dhaliwal, Santosh K. Chaudhary, April Keyes, Ananthan Sadagopan, Lisa Boatner, Neel H. Shah, Charlie Fehl, Keriann M. Backus, Amit Choudhary

**Author notes:** To whom correspondence should be addressed: Amit Choudhary, Chemical Biology and Therapeutics Science, Broad Institute of MIT and Harvard, 415 Main Street, Rm 3012, Cambridge, MA 02142. These authors contributed equally to this work (Endri Karaj, Varsha Venkatarangan, Shaimaa H. Sindi, Surached Siriwongsup, Chaiheon Lee). They will put their name first in their curricula vitae citations or elsewhere. These authors contributed equally to this work (Rajaiah Pergu, Vedagopuram Sreekanth, Karishma Kailass). They will put their name first in their curriculum vitae citations or elsewhere.

## Abstract

Chimeric molecules, which bring together an effector enzyme and a protein-of-interest (POI) to add/remove post-translational modifications (PTMs), are furnishing transformative modalities (e.g., PROTACs). However, these chimeras’ scalability is limited as they employ rare, non-inhibitory binders of effectors. We report GRoup-transfer chimeras for Inducing Proximity (GRIPs) that employ abundantly available effectors’ inhibitors to append POI binder on the effector using group-transfer handles. To demonstrate scalability, we develop 6 GRIPs classes for 3 PTMs utilizing diverse inhibitor, spanning 16 effector‒POI pairs. Furthermore, we report a toolbox of 42 tunable group-transfer handles for Cys/Lys residues and ∼5000 inhibitor‒residue pairs for diverse effectors. Using global proteomics, we confirm the specificity for group transfer and PTM editing. GRIPs endowed new functionalities to POI drugs, including preventing rebound signaling upon drug withdrawal, a more potent/persistent inhibition, and inhibitor-induced pathway activation in 4 fully-endogenous systems. In diverse *hemi*-endogenous systems (tagged POI), GRIPs induced condensate formation with reduced off-targets, cleared pathogenic PTMs, and initiated PTM crosstalk. Overall, GRIPs provide a scalable and versatile platform for PTM editing.

**Graphical abstract:** 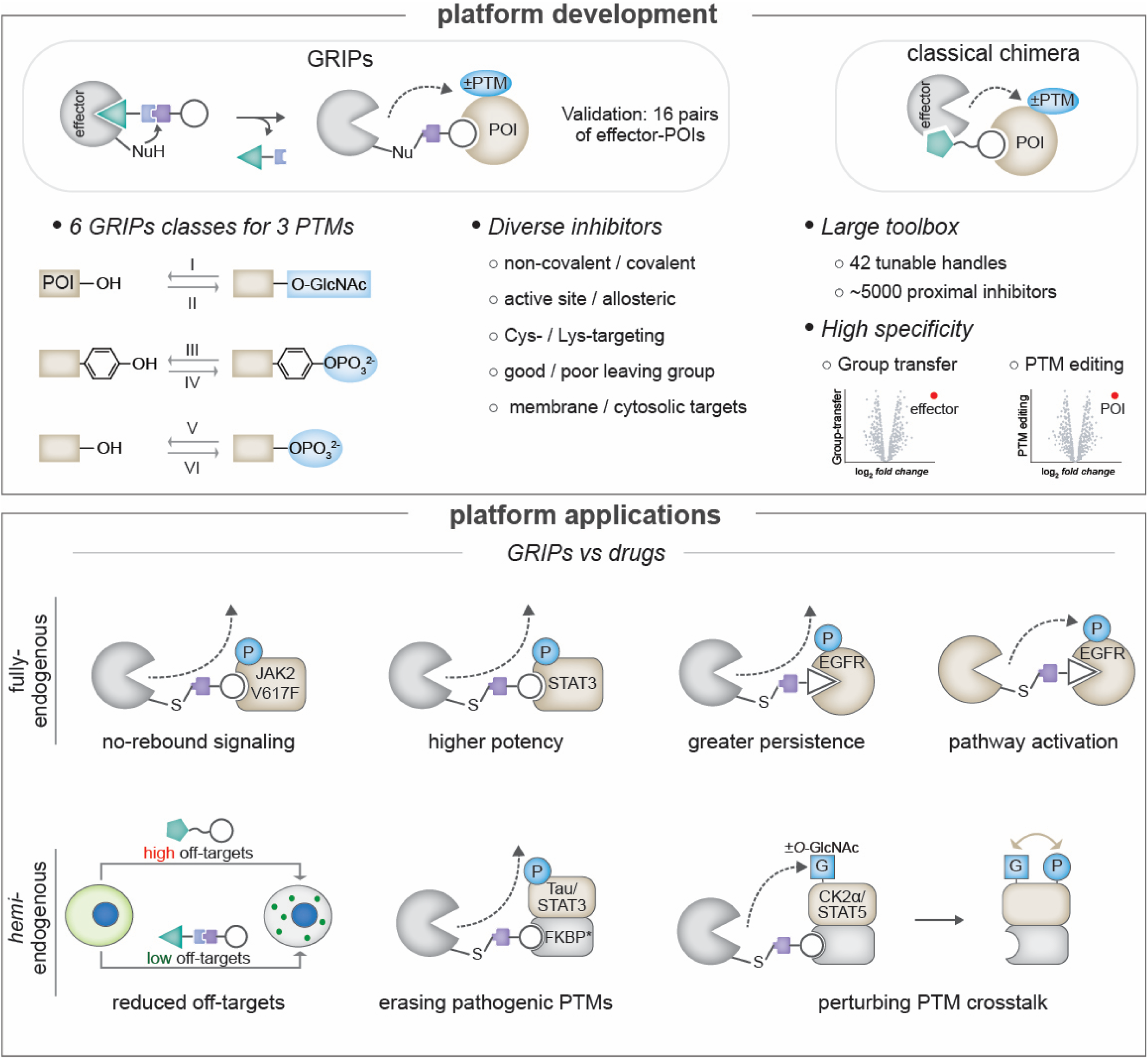

## Introduction

The information flow in biological systems often involves the addition or removal of post-translational modifications (PTMs) by “writer” or “eraser” effector enzymes, with their dysregulation driving numerous pathologies.^1, 2^ Proximity-inducing chimeras are formed by fusing the binders of an effector and a protein of interest (POI). However, as these chimeras must recruit catalytically active writer/eraser, they employ non-inhibitory binders (e.g., activators) that are scarce and challenging to discover, often of poor quality, and some enzymes may not even contain non-inhibitory pockets (Fig. 1A).^3^ Unsurprisingly, such binders are reported for only 3 of 538 human kinases (a well-studied writer),^4–7^ ∼12 of 600 ubiquitin ligases^8–10^ and nearly non-existent for other writers/erasers, restricting chimera development to < 1% of all writers/erasers. A chimera design that allows recruitment of more writers/erasers will be transformative, enabling, for example, PTM editing across diverse sequences since writers/erasers often exhibit sequence specificity.^11–13^

**Figure 1.**
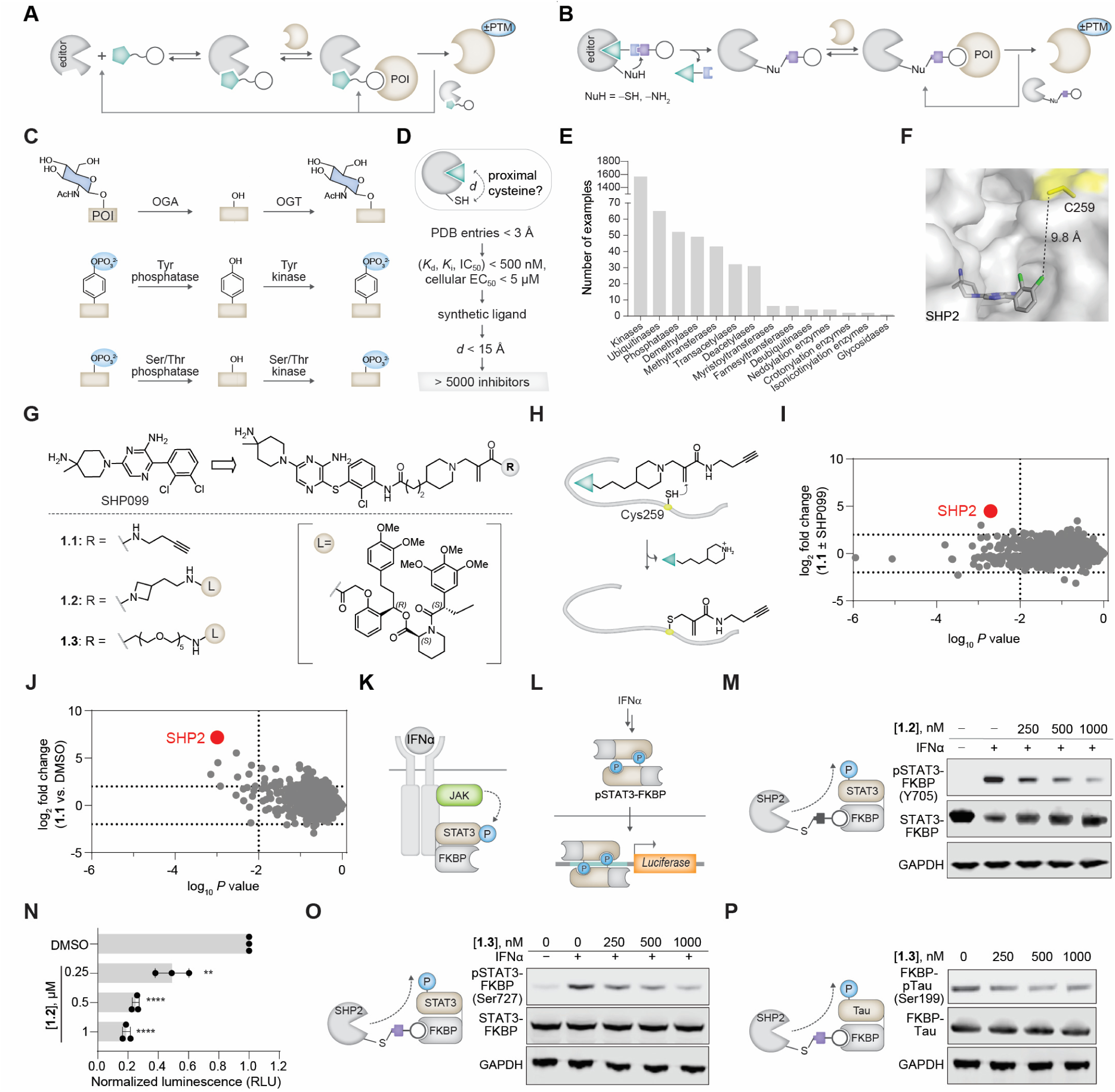
GRIPs concept, bioinformatic studies, and development of dephosphorylation GRIPs. (**A**) Classical chimeras from scarce non-inhibitory binders of effectors. (**B**) A GRIP derived from abundantly available inhibitors. (**C**) *O*-GlcNAc and phosphorylation PTM erasing/writing. (**D**) Identification of proximal cysteine to inhibitor binding pockets using a bioinformatic workflow for PDB data mining. (**E**) Distribution of chemotypes with binding pockets bearing proximal cysteine among different classes of writers/erasers. (**F**) Crystal structure of SHP2 with SHP099 (PDB ID:5EHR) with the targeted cysteine (C259, identified by LC-MS/MS) shown in yellow. (**G**) Structure of GRIPs, alkyne probe **1.1** and FKBP-GRIPs **1.2** and **1.3**. (**H**) A group-transfer reaction to append alkyne on Cys259. (**I,J**) Global proteomic profiling showing labeling of endogenous proteins with alkyne probe **1.1** (100 nM) versus competition with SHP099 (500 nM) (**I**) and DMSO treatment (**J**). Thresholds are defined at fold change [probe/control] = 4 and p value < 0.01. (**K,L**) Schematic of the JAK-STAT pathway (**K**) and luciferase reporter assay (**L**). (**M**) Dose-dependent dephosphorylation of STAT3 (pY705) using GRIP **1.2**. (**N**) Downstream evaluation of STAT3 dephosphorylation using luciferase reporter assay; data present mean values ± SD (n = 3), unpaired *t*-test, two-tailed. STAT3 phosphorylation is induced by IFNα (100 ng/mL). (**O**) Dose-dependent dephosphorylation of STAT3-FKBP (pS727) using GRIP **1.3**. (**P**) Dose-dependent dephosphorylation of Dox-induced FKBP-Tau (pS199) using GRIP **1.3**.

High-quality inhibitors of many writers/erasers are readily available, and a chimera platform from such inhibitors will be scalable (Fig. 1B). Towards that goal, we converted a covalent kinase inhibitor to a chimera that transferred the POI binder onto the kinase’s cysteine using a group-transfer handle.^6^ Under conditions of overexpressed kinase and POI, the chimera induced proximity and triggered POI’s phosphorylation. However, no POI phosphorylation was detectable at the endogenous kinase levels, even with overexpressed POI. Furthermore, to assess the scalability potential and compatibility of the reported group-transfer handles,^6, 14–20^ we generated chimeras from 10 diverse inhibitors (i.e., non-covalent, covalent, allosteric- or active-site-directed), but none induced POI phosphorylation (*vide infra*), even with overexpressed kinases. Thus, we set out to develop group-transfer handles compatible with diverse inhibitors and operative at *endogenous* levels of writer/eraser.

To demonstrate scalability, we set out to develop six classes of chimeras to add or remove three PTMs: *O*-linked *N*-acetylglucosamine (*O*-GlcNAc), Ser/Thr phosphorylation, and Tyr phosphorylation (Fig. 1C).^21–23^ *O*-GlcNAc is appended on Ser/Thr by *O*-GlcNAc transferase (OGT) and erased by *O*-GlcNAcase (OGA).^22^ Ser/Thr and Tyr phosphorylations often employ distinct sets of kinases and phosphatases, differ in their regulatory functions and abundance, prompting calls to classify them as distinct PTMs.^24^ The writers/erasers of these PTMs have readily available inhibitors,^25–30^ and robust assays for PTM detection and downstream signaling, facilitating rigorous validation of the platform. Furthermore, small-molecule chimeras for adding or removing *O*-GlcNAc or Tyr phosphorylation using endogenous writers/erasers (i.e., without overexpression or chemogenetic tagging) do not exist, despite a significant unmet need.^6, 31–34^ While we reported chimeras for endogenous Ser/Thr kinase,^4,7^ they employ activators that induce large off-target phosphorylation from kinase activation (*vide infra*). The reported chimeras from activators of Ser/Thr phosphatase may suffer similarly, in addition to reported concerns of reproducibility and promiscuity^35,36^ in target engagement studies. Thus, these three PTMs provide an opportunity to rigorously validate the platform and ascertain the benefits of a new mode of action.

Herein, we report GRoup-transfer chimeras for Inducing Proximity (GRIPs), which consist of an inhibitor of a writer/eraser connected to the POI binder via a group-transfer handle (Fig. 1B). By mining the Protein Data Bank^37^ and chemoproteomics datasets,^38^ we identified > 5000 inhibitors of diverse writers/erasers with proximal cysteine or lysine. We also developed 42 group-transfer handles with tunable reactivity and length to react with these cysteines or lysines. Using these handles, we report 6 GRIPs classes that recruit *endogenous* writers/erasers to add/remove the 3 PTMs (16 enzyme: POI pairs, 4 fully endogenous), highlighting the platform’s scalability. GRIPs were generated from diverse inhibitors—active-site and allosteric, covalent and non-covalent, and with good or poor leaving groups. The GRIPs platform furnished the first examples of small-molecule chimeras that recruit endogenous writers/erasers of *O*-GlcNAc and tyrosine phosphorylation. GRIPs for Ser/Thr phosphorylation exhibited fewer off-targets than those developed using kinase activators.

Beyond platform development, GRIPs-mediated PTM editing switched on/off multiple biological pathways. Several kinase inhibitor drugs, paradoxically, increase levels of activating phosphorylations, causing life-threatening rebound signaling upon drug withdrawal.^39^ Using Janus Kinase (JAK) drug Baricitinib^40^ and a phosphatase inhibitor, we developed dephosphorylating GRIPs that removed such phosphorylation, preventing rebound signaling of Baricitinib in a cellular model of myeloproliferative neoplasms. Elevated STAT3 phosphorylation is a broad-spectrum cancer driver, but developing effective inhibitors of this transcription factor has been challenging.^41^ In the second application, we report a dephosphorylation GRIP that leverages its “event-driven” pharmacology to attain higher potency in removing STAT3 phosphorylation than a pre-clinical “occupancy-driven” STAT3 inhibitor.^42^ Similarly, dephosphorylation GRIPs based on the EGFR drug Gefitinib afforded a more persistent EGFR inhibition. In a fourth example, we report GRIPs that switch on EGFR signaling using, paradoxically, EGFR inhibitors. These GRIPs mimicked the growth factor EGF used in the biomanufacturing, offering a proteolytically stable, low-cost alternative to EGF. These GRIPs also induced the death of cells harboring oncogenic, but not wild-type KRAS, potentially by perturbing the cancer cells’ goldilocks level of oncogenic signaling.^43^ For the fifth example, we developed GRIPs for dose- and temporal-control of phosphorylation-induced phase separation using tagged POI, achieving similar efficacy to chimeras derived from a kinase activator but with reduced off-targets. Overall, GRIPs provide a scalable approach to add or remove PTMs across diverse biological systems, opening avenues for both basic research and therapeutic development.

## Results

### Inhibitors with proximal cysteines are available for diverse writers and erasers

A scalable GRIPs platform requires readily available inhibitors proximal to a nucleophilic residue to effect a group-transfer reaction. For covalent inhibitors, a proximal nucleophilic residue—typically cysteine—is already available. However, for non-covalent inhibitors, which are more abundant than covalent inhibitors,^44^ a proximal cysteine must be identified. To this end, we queried Protein Data Bank (PDB) to identify human protein: ligand co-crystal structures that meet the following criteria (Fig. 1D): resolution (< 3 Å), biochemical binding constants (*K*_d_, *K*_i_, IC_50_) < 500 nM or cellular EC_50_ < 5 µM, ligand size > 10 carbon atoms, and no cofactors/substrates (e.g., ATP, GTP). Next, we calculated the distance between the ligand and the nearest cysteine on writers or erasers. Since cysteine as far as 15 Å has been targeted by covalent inhibitors,^45^ we chose that cut-off to identify thirteen classes of writers/erasers that have targetable cysteine near the inhibitor (Fig. 1E). A similar search on chemoproteomic datasets of covalent fragments also identified targetable cysteines on many writers and erasers (Extended Data Fig. 1A).^38^ Thus, GRIP-targetable cysteines are available for various writers/erasers and we ventured to develop GRIPs from non-covalent inhibitors, which would be first-in-class.

### Development of a dephosphorylation GRIPs platform that recruits endogenous SHP2 to POI

Our bioinformatic study identified SHP099, an inhibitor of the tyrosine phosphatase SHP2, as proximal to Cys259 (∼10 Å, Fig. 1F).^28^ After confirming that linker attachment to the inhibitor does not perturb its SHP2 binding (*K*_d_ ≈ 30 nM, Extended Data Fig. 1B and Supplementary Fig. 1A),^46, 47^ we synthesized compound **1.1** that will transfer an alkyne group onto SHP2’s Cys259 (Fig. 1G,H), which we confirmed using LC-MS/MS (Fig. MS1). To assess the specificity of group-transfer in cells, we treated HEK293T cells with **1.1**, or a mixture of **1.1** and SHP099 (as a competitor), or DMSO. The lysate was “clicked” with biotin azide, followed by avidin enrichment and a mass spectrometry workflow that allows a pairwise comparison of proteins enriched under probe versus control-treated conditions (Extended Data Fig. 1C). In such pairwise comparisons, SHP2 had the highest labeling fold change (> 64, p value < 0.01) with minimal off-targets (Fig. 1I,J, Supplementary Fig. 1B-F). Interestingly, even those minimal off-targets are unobservable in an equivalent experiment with OGA probe (*vide infra*), suggesting that such off-targets are not inherent to the group-transfer handle.

Next, we developed GRIPs to recruit endogenous SHP2 to a fusion of STAT3 and FKBP12^F36V^ (henceforth called FKBP). In the JAK–STAT pathway, a cytokine (e.g., IFNα) activates JAK, which then phosphorylates and activates the transcription factor STAT (Fig. 1K,L).^48, 49^ We generated a cell line stably expressing FKBP-tagged STAT3, which remained responsive to IFNα stimulation, as evidenced by increased phosphorylation at Y705 (pSTAT3) (Fig. 1M). We synthesized potential SHP2 GRIPs using FKBP ligand (*o*-AP1867)^50^ and various linkers (Fig. 1G and Extended Data Fig. 1D). Upon IFNα stimulation, STAT3 phosphorylation was decreased by several compounds, with **1.2** outperforming those with the PEG linkers (Fig. 1M and Extended Data Fig. 1E,F). GRIP **1.2** triggered a dose-dependent decrease in STAT3 phosphorylation (Fig. 1M), an effect not observed with a mixture of monomers (Extended Data Fig. 1G). To determine the downstream effects of this dephosphorylation, we transfected cells with a STAT3-dependent luciferase reporter (Fig. 1L)^51^ and observed a dose-dependent reduction in reporter activity (Fig. 1N). We note that GRIP **1.2** did not induce dephosphorylation of STAT5, a paralog of STAT3,^52^ but without FKBP fusion (Extended Data Fig. 1H). Global immunoblotting using pan-phosphotyrosine antibody suggested selective dephosphorylation of STAT3‒FKBP (Extended Data Fig. 1I). Collectively, these studies report the first chimeras that recruit endogenous tyrosine phosphatase; these chimeras dephosphorylate STAT3‒FKBP and block the IFNα-triggered JAK-STAT signaling.

While SHP2 is a well-documented tyrosine phosphatase, a study suggested that it can dephosphorylate phospho-serine/-threonine residues.^53^ To determine Ser/Thr dephosphorylation activity of GRIPs, we tested the aforementioned SHP2 GRIPs library on pSer727 on STAT3, which is associated with its oncogenic signaling.^54^ Indeed, GRIPs treatment reduced pSer727 levels but in contrast to tyrosine dephosphorylation, compound **1.3** (with PEG linker) was more effective, suggesting linker-dependent activity (Fig. 1O and Extended Data Fig. 1J-L). Next, we validated this observation on another target: microtubule-associated protein tau (MAPT). Pathological tau phosphorylation (pTau) is a hallmark of neurodegenerative disorders, wherein it reduces tau solubility and microtubule binding while promoting aggregation.^55–59^ We generated a stable cell line with doxycycline-inducible Tau-FKBP and treated the cells with GRIP **1.3**, resulting in a dose-dependent reduction of tau Ser199 phosphorylation,^60^ an early marker of tau pathology (Fig. 1P and Extended Data Fig. 1M-O). Taken together, these studies confirm that SHP2 GRIPs can dephosphorylate serine/threonine residues.

### Fully endogenous dephosphorylation GRIPs inhibit oncogenic JAK-STAT and EGFR signaling

Above, GRIPs were evaluated in *hemi*-endogenous conditions, where the eraser enzyme is endogenous but the POI is tagged. Next, we set out to develop GRIPs for three endogenous POIs—JAK2, STAT3, and EGFR—whose elevated phosphorylation is pathological. For these GRIPs, we use clinical-grade inhibitors of POIs, enabling a direct activity comparisons with GRIPs.

Dysregulated JAKs drive a plethora of pathologies, including 80% of myeloproliferative neoplasms, which frequently harbor the JAK2^V617F^ gain-of-function variant, leading to persistent JAK-STAT signaling.^61–65^ Several type I ATP-competitive JAK2 inhibitors have emerged, but they paradoxically increase the levels of JAK2-activating phosphorylation.^66, 67^ Here, the inhibitor stabilizes the JAK2 conformation, restricting phosphatases’ access to the activating phosphorylation.^68^ The result is the accumulation of hyperphosphorylated JAK2 (Fig. 2A) that is “primed” for signaling upon inhibitor withdrawal.^69^ Clinically, this withdrawal triggers respiratory failure and septic shock-like syndrome,^70, 71^ and is associated with cardiotoxicity of JAK inhibitors.^72^

**Figure 2.**
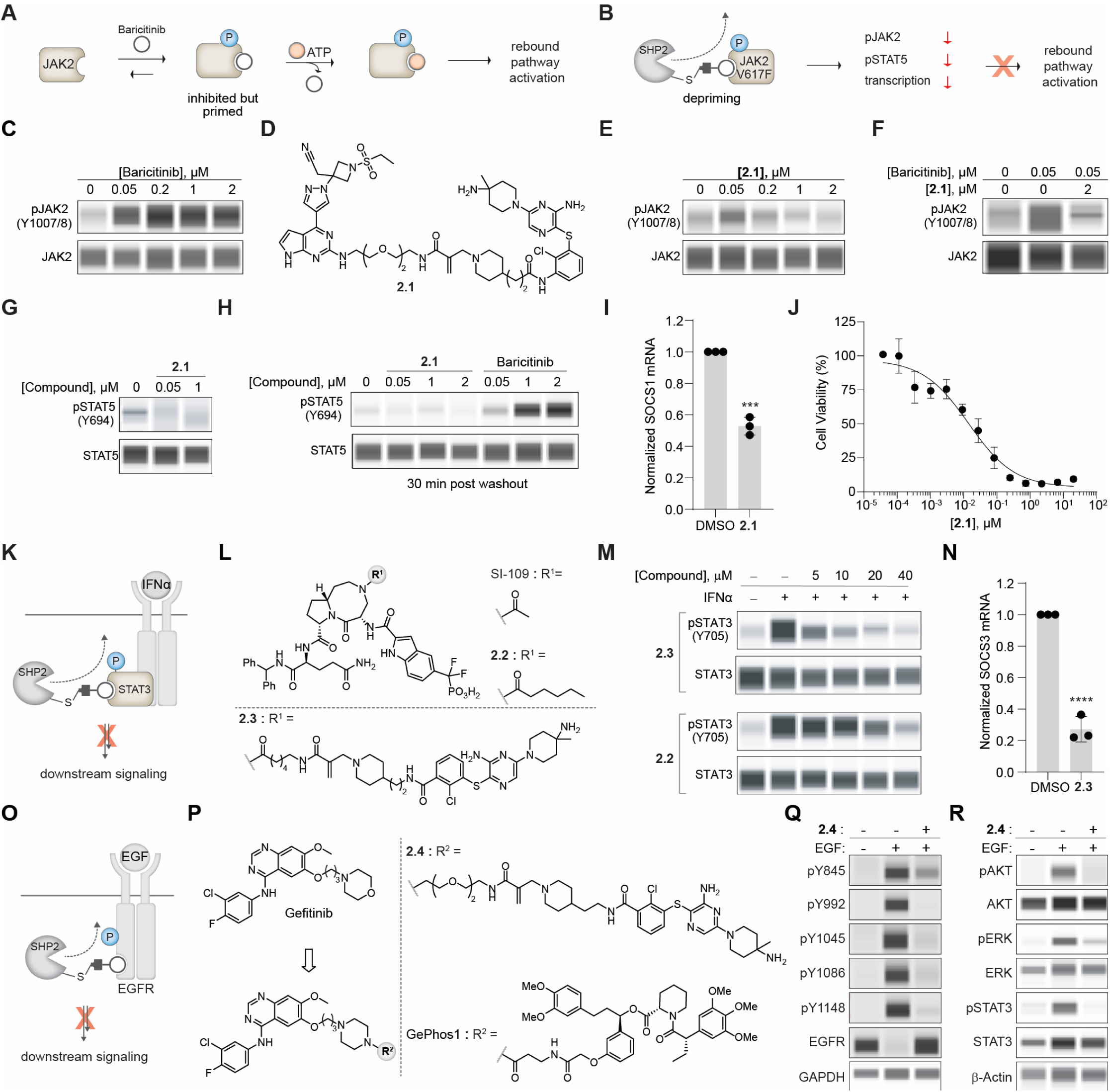
Fully endogenous applications of dephosphorylation GRIPs. (**A**) Schematic of paradoxical JAK2 activation upon drug removal. (**B**) Effect of SHP2‒JAK2^V617F^ GRIPs. (**C**) Phosphorylation of JAK2 after treatment with Baricitinib (0.05–2 μM). (**D**) Structure of GRIP **2.1**. (**E**) Phospho- (Y1007/8) levels of JAK2 after treatment with GRIP **2.1**. (**F**) Baricitinib competition assay with GRIP **2.1**. (**G**) Phospho-STAT5 (Y694) after treatment with GRIP **2.1**. (**H**) pSTAT5 levels of HEL92.1.7 cells treated with Baricitinib or **2.1** followed by washout. (**I**) Quantitation of SOCS1 mRNA using a qPCR assay for DMSO or GRIP **2.1** (5 μM)-treated cells. Data present mean values ± SD (n = 3), unpaired *t*-test, two-tailed. (**J**) Cytotoxicity assay of **2.1** in SET2 cells. (**K**) Schematic of endogenous STAT3 dephosphorylation by GRIPs. (**L**) Structures of STAT3 inhibitor SI-109, control inhibitor **2.2**, and GRIP **2.3**. (**M**) Dose-dependent dephosphorylation of STAT3 (pY705) using GRIP **2.3** and control inhibitor **2.2**. (**N**) Quantitation of SOCS3 mRNA using a qPCR assay for DMSO or GRIP **2.3** (5 μM)-treated cells. Data present mean values ± SD (n = 3), unpaired *t*-test, two-tailed. For all STAT3 dephosphorylation assays, STAT3 phosphorylation is induced by IFNα (100 ng/mL). (**O**) Schematic of endogenous EGFR dephosphorylation by GRIPs. (**P**) Structure of SHP2‒EGFR GRIP **2.4** and control compound GePhos1. (**Q**) Dephosphorylation of EGFR at various phosphosites by GRIP **2.4** (250 nM). (**R**) Assessment of the phosphorylation status of AKT, ERK, and STAT3 upon GRIP **2.4** (250 nM) treatment.

We hypothesized that SHP2 GRIPs from these inhibitors could remove the activating phosphorylation and rebound signaling (Fig. 2B). We treated HEL 92.1.7 erythroleukemia cells^62^ (expressing JAK2^V617F^) with JAK inhibitor Baricitinib and confirmed the elevation of activating phosphorylation on JAK2 (Fig. 2C).^69, 72^ We used Baricitinib to generate the SHP2‒JAK2 GRIP **2.1** (Fig. 2D) and treated HEL92.1.7 cells with increasing doses of **2.1**. At a lower dose (50 nM), we observed an increase in pJAK2, indicating target engagement. However, at higher doses, we observed a decrease in pJAK2, suggesting SHP2 recruitment that removes the phosphorylation (Fig. 2E). We conducted a competition experiment in which we treated HEL92.1.7 cells with Baricitinib (50 nM) or Baricitinib and **2.1** (2 μM) and observed a decreased JAK2 phosphorylation in the latter condition (Fig. 2F). As expected, GRIPs-induced JAK2 dephosphorylation resulted in decreased phospho-STAT5 levels, consistent with inhibition of downstream signaling (Fig. 2G).

To assess the impact of rebound signaling upon Baricitinib withdrawal, we treated HEL92.1.7 cells with either GRIPs or Baricitinib, and then determined pSTAT5 levels after a washout. Consistent with previous reports,^69, 72^ Baricitinib-treated cells exhibited elevated pSTAT5 levels compared to DMSO controls, reflecting rapid reactivation of signaling from a primed pool of phosphorylated JAK2 (Fig. 2H). In contrast, GRIPs-treated cells displayed pSTAT5 levels comparable to those of DMSO-treated cells, indicating effective JAK2 depriming and absence of rebound signaling (Fig. 2H). We also performed qPCR to quantify SOCS1 mRNA levels, a downstream effector gene in the pathway.^73^ Treatment with GRIP **2.1** resulted in a significant reduction in SOCS1 levels compared with DMSO-treated cells (Fig. 2I). Furthermore, we observed potent and dose-dependent cytotoxic effects in a cancer cell line model used for assessment of JAK2 inhibitors (Fig. 2J). Overall, these studies show that SHP2‒JAK2 GRIP can remove the rebound signaling side effect of JAK2 inhibitors, offering an induced proximity strategy to overcome the limitations of occupancy-based inhibitors.

In a broad spectrum of cancers, STAT3 is persistently active, driving tumor growth and metastasis^48, 49^ and a dephosphorylating GRIPs could potentially shut down such tumorigenic signaling (Fig. 2K). Towards that goal, we replaced the FKBP binder in GRIP **1.2** with the selective STAT3 inhibitor SI-109 to generate a potential SHP2‒STAT3 GRIP **2.3** (Fig. 2L) and used a derivatized analog of SI-109 (**2.2**) as a control.^74^ For synthetic convenience, we reversed the amide connectivity of the group-transfer handle attached to the SHP2 inhibitor and confirmed cellular target engagement as done above (Extended Data Fig. 2A-C and Supplementary Fig. 1G). We treated HEK293T cells with IFNα and either **2.3** or the control inhibitor **2.2**, and observed a dose-dependent dephosphorylation of endogenous STAT3 upon GRIPs treatment, an effect not observed with the control inhibitor **2.2**, which required a concentration >4 times higher to achieve the same effect (Fig. 2M). We also performed a qPCR experiment to determine SOCS3 mRNA levels, a direct STAT3 transcriptional target.^73^ Treatment with **2.3** significantly reduced SOCS3 mRNA levels compared with DMSO-treated cells (Fig. 2N). These studies report a more potent approach to inhibit STAT3 than that afforded by an “occupancy-based” inhibitor, and a distinct mechanism of action from that of occupancy-based inhibitors or PROTACs.^74, 75^

Our third example of endogenous dephosphorylation GRIPs was inspired by the work of Crews and co-workers, who reported a dual-inhibitory approach to EGFR. Here, an engineered phosphatase (FKBP-PTPN2) was recruited to EGFR using a chimera (termed GePhos1) composed of the FKBP binder and the EGFR inhibitor Gefitinib.^76^ This study suggested that the dual inhibition approach will be more potent inhibition with a differential downstream gene expression than Gefitinib and has the potential to overcome EGFR drug resistance. Motivated by these findings, we designed SHP2‒EGFR GRIPs **2.4** (Fig. 2O,P). Following the reported protocol,^76^ we treated HeLa cells with GRIPs **2.4** or GePhos1 and stimulated cells with EGF. GRIPs **2.4** treatment reduced phosphorylation at the key Y845 residue and at other sites (Fig. 2Q), and this dephosphorylation diminished phosphorylation of downstream effectors, including AKT, ERK, and STAT3 (Fig. 2R). When comparing the effects of GRIP **2.4** to GePhos1, we observed differential and enhanced dephosphorylation, which was associated with more potent downstream inhibition (Extended Data Fig. 2D,E).^76^ Importantly, in a washout experiment involving treatment of Hela cells with **2.4** or GePhos1 followed by washout and monitoring of pEGFR levels, **2.4** displayed higher reduction in phosphorylation levels at multiple phosphosites compared with that of GePhos1-treated cells, suggesting that GRIPs confers a more sustained inhibition than occupancy-based inhibitors (Extended Data Fig. 2F).^76–80^ In summary, we report 3 GRIPs that provide new attributes to known drugs.

### Development of a de-*O*-GlcNAcylation GRIPs platform that recruits endogenous OGA to POIs

The workflow for development of OGA GRIPs mirrored those of SHP2, starting with bioinformatic search that identified OGA inhibitor Thiamet-G to be proximal to cysteine 316 (8 Å, Fig. 3A). Importantly, Thiamet-G analogs with diverse binding affinities are available,^81^ allowing assessment of the impact of ligand affinity on group-transfer efficacy and specificity (Extended Data Fig. 3A). Guided by the co-crystal structure,^82^ we synthesized compounds **3.1** and **E3.4** by connecting a potent inhibitor (*K*_i_ = 2.1 nM) or a weak inhibitor (*K*_i_ ≈ 1 µM)^81^ to a group-transfer handle with alkyne (Fig. 3B, Extended Data Fig. 3A). Upon binding, these compounds should transfer the alkyne group to OGA’s cysteine residue, which can be detected by mass spectrometry, enabling assessment of both group-transfer efficacy on OGA and specificity. Treatment of HEK293T cells with **3.1** and pairwise comparison with competition or DMSO suggested remarkable selectivity for OGA, with **3.1** showing an enrichment >128-fold relative to DMSO-treated samples (Fig. 3C,D). Alkyne probe **E3.4,** based on a weaker inhibitor, while still engaging OGA at a higher concentration, exhibited several off-targets (Extended Data Fig. 3B) and was unable to sufficiently label OGA even when overexpressed (Extended Data Fig. 3C). In contrast, **3.1** could label both overexpressed and endogenous OGA with high selectivity (Extended Data Fig. 3C,D). Further LC-MS/MS studies confirmed Cys316 as the primary labeled residue (Fig. MS2).

**Figure 3.**
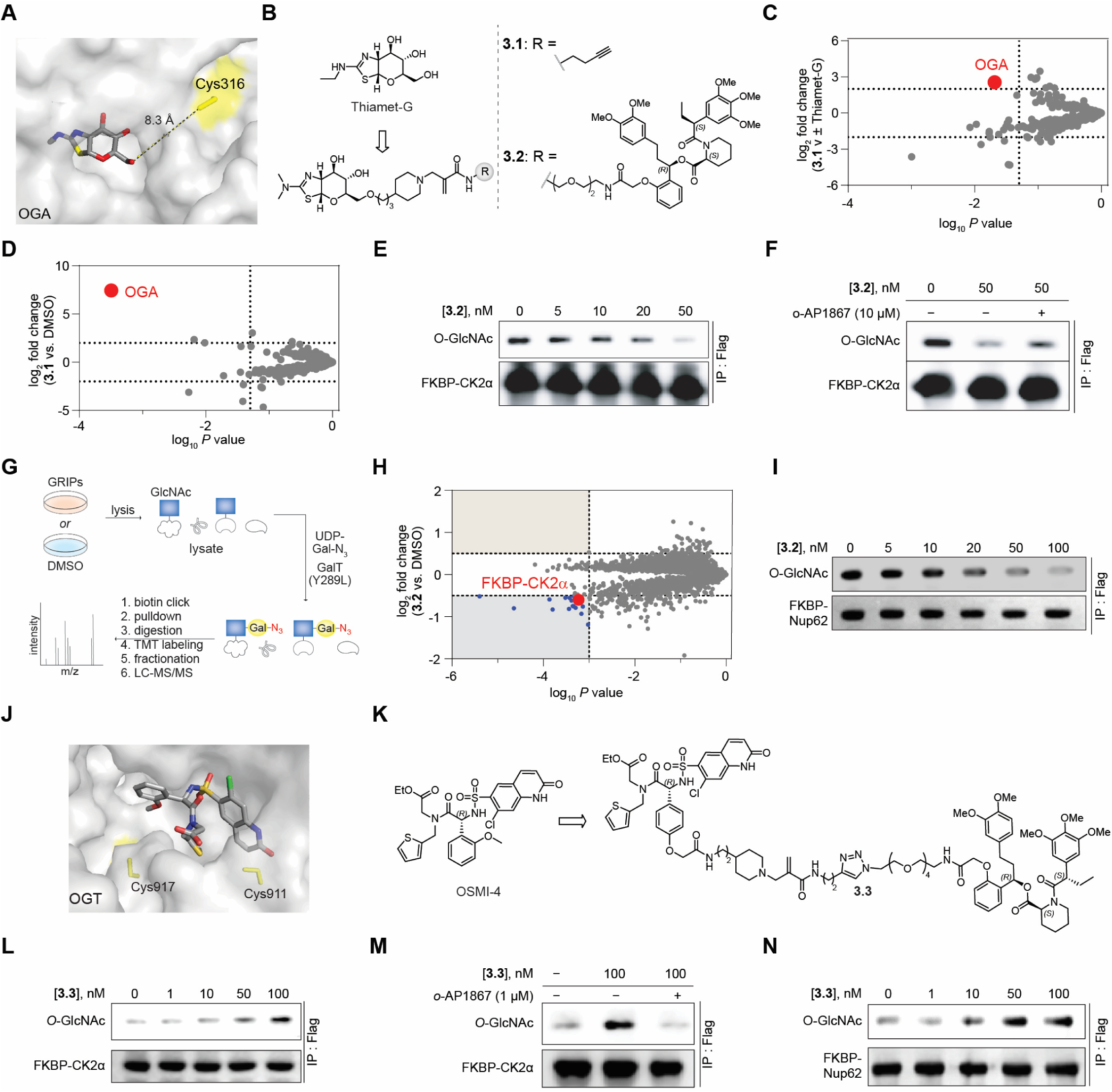
GRIPs for adding and removing *O-*GlcNAc. (**A**) Crystal structure of OGA with Thiamet-G (PDBid: 5M7S) with the targeted cysteine (C316, identified by LC-MS/MS) highlighted in yellow. (**B**) Structures of Thiamet-G and analogs, which are derivatized with an alkyne click handle **3.1** and FKBP binder **3.2**. (**C,D**) Global proteomic profiling showing labeling of endogenous proteins with GRIP **3.1** (100 nM) with Thiamet-G competition (**C**) or versus DMSO (**D**). Thresholds are defined at fold change [probe/control] = 4 and p value < 0.05. (**E**) Dose-dependent deglycosylation of FKBP-CK2α using GRIP **3.2**. (**F**) Competition-dependent deglycosylation of FKBP-CK2α using GRIP **3.2.** (**G**) Schematic of workflow for global quantitative *O*-GlcNAc proteomic profiling. (**H**) Global *O*-GlcNAc proteomic profiling of GRIP **3.2** (50 nM) using DMSO treated HEK293T cells (transiently expressing FKBP-CK2α) as control. Thresholds are defined at fold change [GRIP/control] = 1.4 and *p* value < 0.001. (**I**) Dose-dependent deglycosylation of FKBP-Nup62 using GRIP **3.2**. (**J**) Crystal structure of OGT with OSMI-4 (PDBid: 6MA1) and one of the labeled cysteines (C911, identified by LC-MS/MS) is highlighted in yellow. (**K**) Structure of OSMI-4 and the corresponding GRIP **3.3**. (**L**) Dose-dependent glycosylation of FKBP-CK2α using **3.3**. (**M**) Competition of **3.3** effects by FKBP binder, *o*-AP1867. (**N**) Dose-dependent glycosylation of FKBP-Nup62 using **3.3**.

Overall, target engagement studies for SHP2 and OGA, and reported group-transfer handle for chemogenetic tags,^83^ suggest high specificity of the group-transfer handle.

Since OGA is known to remove *O*-GlcNAc from casein kinase 2α (CK2α),^84^ we sought to develop OGA GRIPs targeting an FKBP-CK2α fusion. We synthesized potential OGA GRIPs using FKBP ligand and linkers of different lengths and hydrophobicity (Fig. 3B and Extended Data Fig. 3E). We rapidly screened our GRIPs library using a co-transfection system (FKBP-CK2α:OGT:OGA in 4:3:1 ratio) that enhances the *O*-GlcNAc signal in an assay involving immunoprecipitating CK2α and immunoblotting for *O*-GlcNAc. These screening efforts revealed dose-, binder-, and linker-dependent removal of *O*-GlcNAc from CK2α, with PEG linkers being more efficacious than alkyl-based linkers. Notably, GRIPs based on the weaker OGA inhibitor could still induce *O*-GlcNAc removal, but at higher concentrations (Extended Data Fig. 3F-I).

Next, to determine if GRIP **3.2** can recruit endogenous OGA to remove *O*-GlcNAc from FKBP-CK2α, we treated HEK293T cells transiently expressing FKBP-CK2α with **3.2**. Subsequent immunoprecipitation and immunoblotting for *O*-GlcNAc showed a dose-dependent deglycosylation of FKBP-CK2α (Fig. 3E), an effect that was abolished by excess FKBP ligand (Fig. 3F). To assess the proteome-wide selectivity of *O*-GlcNAc removal by **3.2**, we performed global proteomics. HEK293T cells were treated with DMSO or **3.2**, followed by chemoenzymatic labeling of *O*-GlcNAc–modified proteins using GalT(Y289L) and GalNAc-azide (Fig. 3G).^85^ Labeled proteomes were analyzed using a quantitative LC–MS/MS workflow (Fig. 3G), revealing a significant reduction in *O*-GlcNAc levels on FKBP–CK2α in GRIP-treated cells relative to DMSO controls (Fig. 3H). In addition, a limited set of other proteins exhibited reduced *O*-GlcNAcylation, many of which are known substrates or interactors of CK2α substrates (e.g., ELF1, RBPJ, MEF2D).^86^ As *O*-GlcNAcylation of CK2α modulates its kinase activity,^84^ we expect this *O*-GlcNAc removal to perturb downstream effectors,^84^ particularly given the competition between phosphorylation and *O*-GlcNAcylation.^87, 88^ Notably, no proteins displayed increased *O*-GlcNAcylation, indicating that **3.2** does not globally inhibit endogenous OGA activity (Fig. 3H). In addition to FKBP–CK2α, we confirmed *O*-GlcNAc removal on another protein, Nup62, a nucleoporin known to be highly *O*-GlcNAcylated (Fig. 3I).^89^ Analogous to the FKBP-CK2α experiments, FKBP-Nup62 was transiently expressed in HEK293T cells and incubated with GRIP **3.2** to yield a dose-dependent removal of *O*-GlcNAc. Collectively, these studies report the first examples of small-molecule chimeras that recruit endogenous OGA to remove *O*-GlcNAc from a tagged POI and we confirm the specificity of such removal using global proteomics.

### Development of an *O*-GlcNAcylation GRIPs platform that recruits endogenous OGT to POIs

To complement OGA GRIPs, we developed OGT GRIPs using OSMI-4, which has high biochemical potency albeit suboptimal cellular efficacy (Fig. 3J,K).^23, 29^ Guided by the co-crystal structure (Fig. 3J),^29^ we identified two exit vectors (on the phenyl ring or near amide nitrogen) to append a group-transfer handle with alkyne, yielding compounds **E3.13** and **E3.14** (Extended Data Fig. 3J). Both compounds labeled transiently expressed OGT in HEK293T cells (Extended Data Fig. 3K), thereby validating both exit vector strategies, but we prioritized the **E3.13** inhibitor scaffold owing to synthetic convenience. Treatment of recombinant OGT with **E3.13** and subsequent LC-MS/MS studies revealed labeling at Cys911, a cysteine within the binding pocket and some labeling at a more distal cysteines Cys417 and Cys531 (Fig. MS3). We next synthesized potential OGT GRIPs **3.3** to install *O*-GlcNAc on FKBP-CK2α (Fig. 3K). As above, we transfected HEK293T cells with FKBP-CK2α and observed a dose-dependent induction of *O*-GlcNAcylation on FKBP-CK2α (Fig. 3L) and a loss of O-GlcNAcylation when the FKBP binder was used as a competitor (Fig. 3M). As with OGA GRIPs, we confirmed that GRIPs **3.3** could induce *O*-GlcNAcylation of Nup62 (Fig. 3N).

We also confirmed previous reports on crosstalk between *O*-GlcNAcylation and phosphorylation on STAT5. Briefly, removal of *O*-GlcNAc from STAT5 destabilizes its conformation, preventing its binding to JAK2 and inhibiting phosphorylation (Supplementary Fig. 2A).^90^ In a cell line stably expressing FKBP-STAT5, OGA GRIPs **1.2** induced a dose-dependent reduction in levels of phosphorylation on STAT5, and this loss of phosphorylation was recovered when the FKBP binder *o*-AP1867 was used as a competitor (Supplementary Fig. 2B). Furthermore, OGT GRIPs **3.3** induced a dose-dependent increase in STAT5 phosphorylation (Supplementary Fig. 2C,D). Taken together, these studies describe the first examples of endogenous OGA or OGT recruitment to remove or add *O*-GlcNAc from FKBP-tagged POIs, providing tools to study emerging *O*-GlcNAc biology.

### Contemporary group-transfer handle cannot furnish GRIPs from common covalent inhibitor scaffolds

With the successful development of GRIPs from non-covalent inhibitors, we sought to scale up the GRIPs platform to cysteine-based covalent inhibitors. These inhibitors contain a reactive group (e.g., a Michael acceptor) that can be replaced by a group-transfer handle, with the connecting nitrogen functioning as the departing moiety in the group-transfer reaction (Fig. 4A).^6^ While we have successfully developed GRIPs-like chimeras by such a swap, we reasoned this approach is not scalable (Fig. 4A).^6^ We hypothesized that while the methacrylamide handle exhibits optimum reactivity to cysteine when the covalent inhibitor has a strongly basic nitrogen (Fig. 4B),^6, 14–16^ most covalent inhibitors bear weakly basic nitrogen (Fig. 4C) for which the reactivity should be poor. The strongly basic nitrogen is protonated under physiological conditions, which enhances group-transfer reactivity. To confirm this hypothesis, we chose four covalent inhibitors with weakly basic nitrogen and appended the reported methacrylamide handle (Fig. 4C, Supplementary Fig. 2E). Unfortunately, none of these compounds underwent group- transfer reaction with cysteine under biochemical conditions (Supplementary Fig. 2F). To confirm activity in cells, we appended bromodomain-containing protein 4 (BRD4)^91^ binder (*S*)-JQ1 to these compounds to generate potential GRIPs that would induce proximity between BRD4 and kinase, resulting in BRD4 phosphorylation (Supplementary Fig. 2E). We measured BRD4 phosphorylation using a previously reported assay^6^ involving immunoblotting of immunoprecipitated BRD4 in HEK293T cells transiently expressing the kinase and BRD4. Consistent with the observed lack of cysteine reactivity, we did not detect BRD4 phosphorylation in compound-treated cells (Supplementary Fig. 2G-J). Thus, scaling up the GRIPs platform to all covalent inhibitors requires group-transfer handles compatible with weakly and strongly basic nitrogen.

**Figure 4.**
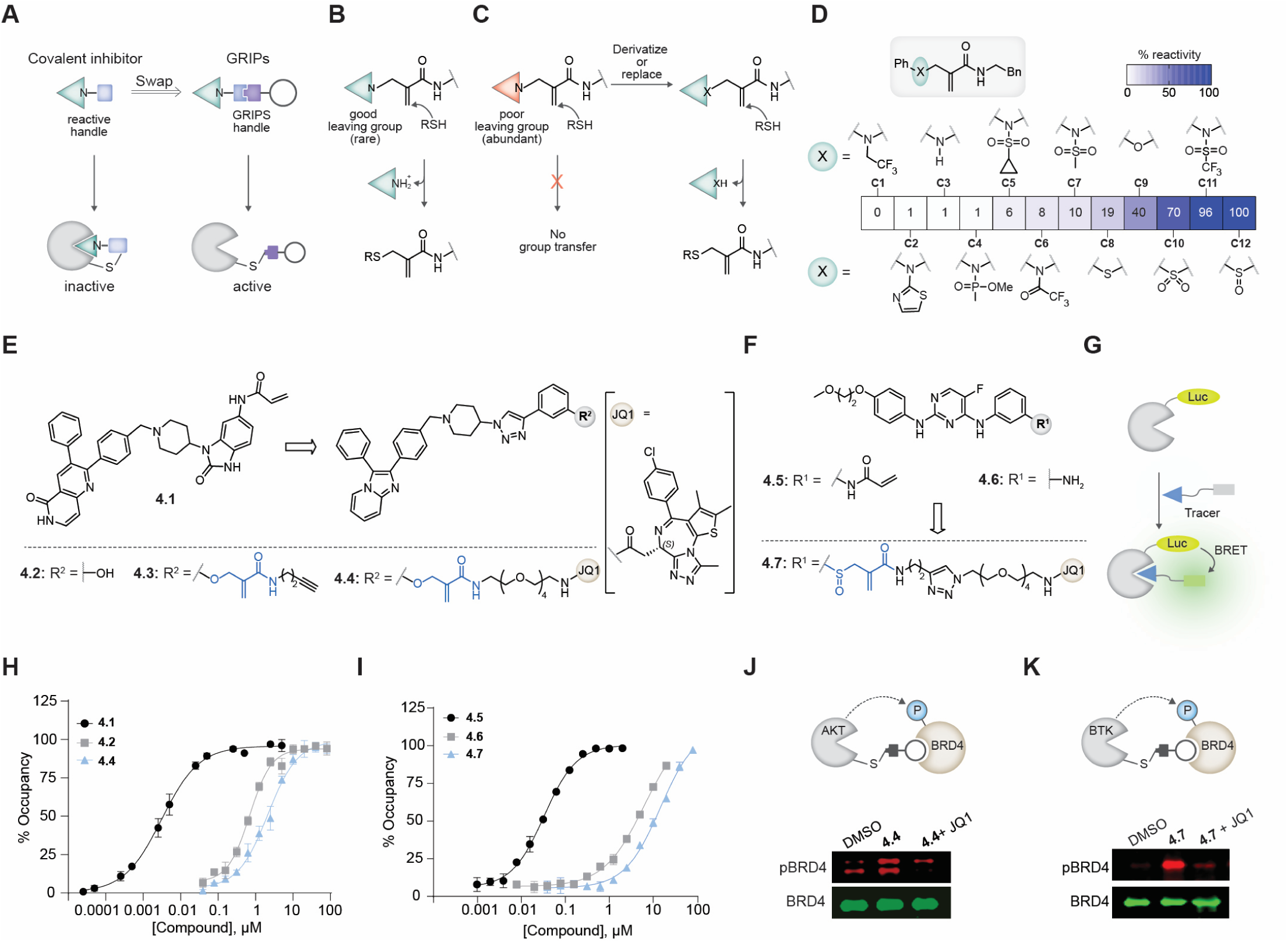
GRIPs from covalent inhibitors with poor leaving groups (weakly basic nitrogen). **(A)** Development of GRIPs from covalent inhibitors. **(B)** Covalent inhibitors with good leaving groups are less abundant. (**C**) Poor leaving group ability can inhibit group-transfer reactivity. Derivatizing or replacing poor leaving groups is expected to generate functional group-transfer handles. (**D**) Structures and percent reactivity of model compounds with *N*-acetyl-cysteine-OMe normalized against the least and most reactive compound. The heat map represents % cysteine reactivity (mean value, of n = 3). (**E,F**) Structures of AKT covalent inhibitor Borussertib **4.1**, non-covalent phenol **4.2**, GRIP alkyne **4.3** and BRD4-GRIP **4.4** (**E**) and of BTK covalent inhibitor Spebrutinib **4.5**, non-covalent amine **4.6**, and BRD4-GRIP **4.7** (**F**). (**G**) Schematic of nanoBRET assay. (**H,I**) ATP pocket occupancy of covalent inhibitors, non-covalent binders, and GRIPs targeting AKT (**H**) or BTK (**I**). Data are presented as mean values ± SD (n = 2). (**J,K**) BRD4 phosphorylation by AKT GRIP **4.4** (**J**, 250 nM) or BTK GRIP **4.7** (**K**, 250 nM) with competition with (*S*)-JQ1 (1 μM).

### Development of GRIPs compatible group-transfer handles for common covalent inhibitor scaffolds

To develop GRIPs from covalent inhibitors with weakly basic nitrogen, we generated a set of tunable handles by either late-stage functionalization of nitrogen or by replacing it with oxygen or sulfur as a minimalistic change (Fig. 4D). For functionalization, we appended an electron-withdrawing –CF_3_ (**C1**) or an aminothiazole (**C2**) moiety, generating sulfonamides (**C5**, **C7**, and **C11**), phosphoramides (**C4**), and acetamides (**C6**), thereby expanding the reactivity range. Replacement with oxygen (**C9**) or sulfur (**C8**), the latter of which was oxidized to sulfoxide (**C12**) or sulfone (**C10**), enabled further fine-tuning of reactivity. These handle designs mimic the electronic effects of protonation by attenuating the nitrogen lone-pair resonance with the π system or by increasing electron-withdrawal. Such mimicry enabled the development of a suite of minimalistic group-transfer handles with tunable reactivity, 6‒100-fold greater than that of unreactive nitrogen.

With these handles in hand, we generated potential BRD4 GRIPs from the allosteric AKT inhibitor Borussertib (**4.1**, Fig. 4E)^30^ and the ATP-competitive Bruton’s tyrosine kinase (BTK) inhibitor Spebrutinib^27^ (**4.5**, Fig. 4F), both of which have an unreactive nitrogen. As done with non-covalent inhibitors, we generated alkyne probes for AKT **4.3** and **E4.1** (Fig. 4E and Extended Data Fig. 4A) and confirmed the target engagement and specificity using global proteomics (Extended Data Fig. 4B-D). Furthermore, using LC-MS/MS, we confirmed that the alkyne probe labeled the identical cysteine as the parent covalent inhibitor (Extended Data Fig. 4E, Supplementary Fig. 2K,L, and Fig. MS4). Using a BRET assay, we confirmed the release of the inhibitor from the ATP pocket following a group-transfer reaction of GRIPs.^92, 93^ Here, BRET arising from the tracer that binds to the ATP pocket and the AKT or BTK-fused nanoluciferase reports on ATP pocket occupancy (Fig. 4G). HEK293T cells transiently expressing AKT or BTK-nanoluciferase were treated with covalent inhibitors Borussertib **4.1** and Spebrutinib **4.5**, non-covalent inhibitors (**4.2** and **4.6**), or BRD4 GRIPs (**4.4** and **4.7**). While covalent inhibitors decreased BRET, the GRIPs or non-covalent inhibitors did not, confirming that the kinase is not inhibited by the GRIPs (Fig. 4H,I, and Extended Data Fig. 4 F,G). Congruently, BRD4 phosphorylation was observed with GRIPs with a reactive handle (e.g., phenol or sulfoxide) but not with compounds with poor reactivity (Fig. 4J,K, and Extended Data Fig. 4H). Furthermore, an excess of the BRD4 binder (*S*)-JQ1 acted as a competitor, reducing BRD4 phosphorylation (Fig. 4J,K). Overall, we report tunable group-transfer handles compatible with inhibitors bearing both good and poor leaving groups, enabling the development of GRIPs from a wide range of covalent inhibitors.

### Development of group-transfer handles and GRIPs platform with lysine as the nucleophile

Many writers/erasers may lack a targetable cysteine, prompting us to explore if lysine can be the nucleophile, given its greater abundance relative to cysteine (Fig. 5A).^94^ Indeed, an examination of protein: ligand structures in the Protein Data Bank (PDB) suggests that ≈97% of proteins have a lysine residue within 15 Å (Supplementary Fig. 3A).^94^ Using the aforementioned bioinformatic workflow (Fig. 1D), we identified inhibitors of several writers/erasers with proximal lysine (Fig. 5B).

**Figure 5.**
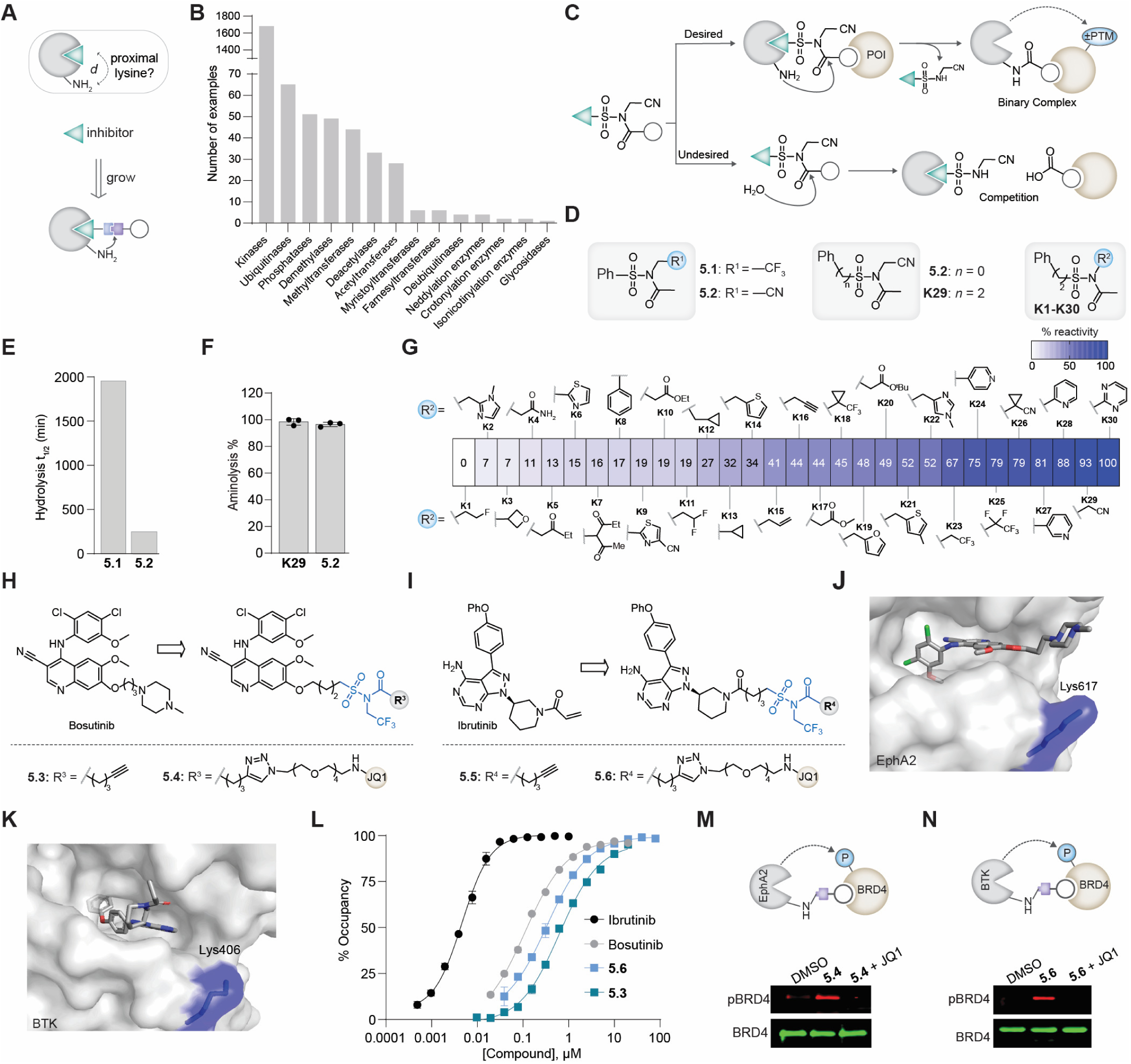
Lysine-targeting GRIPs from non-covalent inhibitors. (**A**) Schematic of the conversion of non-covalent inhibitors to GRIPs using the lysine group-transfer. (**B**) Available inhibitors of diverse writers/erasers proximal to a lysine. (**C**) Productive and unproductive pathways for NASA-GRIPs. (**D**) Structures of NASA and SuFA fragments. (**E**) Replacement of nitrile group (**5.2**) in NASA with trifluoromethyl (**5.1**) increases hydrolytic stability. (**F**) Size reduction of NASA by replacing phenyl (**5.2**) with methylene (**K29**) and assessment of their corresponding lysine reactivity. Compounds **5.2** and **K29** were reacted with *N*-Boc-lysine-coumarin, and the corresponding acylated product was quantified by LC-MS analysis. Data are presented as mean values ± SD, n = 3. (**G**) A toolbox of group-transfer handles with tunable reactivity to lysine. The heat map represents normalized lysine reactivity (0%: least reactive, 100 %: most reactive). (**H,I**) Structures of lysine-based GRIPs **5.3-6** targeting EphA2 (**H**) and BTK (**I**). (**J**) Alkyne GRIP **5.3** labels EphA2 at Lys617. Crystal structure obtained from PDB ID: 5I9X. (**K**) Alkyne **5.5** labels BTK at Lys406. Crystal structure obtained from PDB ID: 5P9J. (**L**) Nano-BRET assay for Ibrutinib, Bosutinib, **5.3** and **5.6**. Data are presented as mean ± SD, n = 2 **(M,N)** GRIPs **5.4** and **5.6** targeting EphA2 (**M**) and BTK (**N**), respectively, induced BRD4 phosphorylation, which is competed out by excess (*S*)-JQ1. Compounds **5.4** and **5.6** were evaluated in transfected HEK293T cells expressing both kinase and BRD4 and treated with 250 nM of GRIPs (**5.4** and **5.6**) and for competition with 1 μM (*S*)-JQ1

We initiated our studies on developing lysine-targeting GRIPs by examining extant lysine-based group-transfer handles based on *N*-acyl-*N*-alkyl phenylsulfonamides (NASA).^17–20^ Briefly, we generated potential BRD4-targeting GRIPs from six kinase inhibitors and examined their ability to induce BRD4 phosphorylation. As with efforts above with extant cysteine group-transfer handles, none of the NASA-based compounds induced BRD4 phosphorylation (Supplementary Fig. 3B-O). We attribute this failure to multiple reasons. First, NASA is hydrolytically unstable (*t*_1/2_ ∼ 5 h in PBS) and has a short shelf life, making working with these reagents challenging.^19^ Furthermore, the hydrolytic instability can decrease GRIPs’ activity by generating monomeric components (Fig. 5C) that compete with GRIP for POI and enzyme binding. This competition can be significant, as the working concentration of most GRIPs is in the low nanomolar range. Furthermore, NASA’s high non-specific labeling, which is well-documented by multiple groups,^19, 95, 96^ will have effects akin to hydrolysis, reducing its suitability for induced proximity applications.^31, 97^ Second, the aryl group in NASA contributes to steric hindrance and handle rigidity in GRIPs and may impede access to lysine in deep pockets.

Finally, since the pK_a_ (and reactivity) of lysine varies significantly on enzymes,^98^ group-transfer handles with tunable reactivity will be needed to generate specific GRIPs. Thus, as described above for cysteine group-transfer handles, we developed a set of lysine group-transfer handles via systematic chemical modifications of NASA to enhance hydrolytic stability, size reduction, and increase tunability (Fig. 5D).

Studies on carboxypeptidases^99^ and activated amides^100, 101^ have shown that hydrogen bond acceptors (e.g., nitrile in NASA) can facilitate amide hydrolysis. We envisioned that the replacement of the nitrile with a trifluoromethyl group (─CF_3_) would enhance hydrolytic stability, as ─CF_3_ is a poor hydrogen bond acceptor^102^ while still being a potent electron-withdrawing group. Indeed, such replacement increased hydrolytic stability by ∼5-fold (Fig. 5D,E). Furthermore, unlike the phenyl group in benzamide, the phenyl group in benzenesulfonamide is not in strong resonance;^103^ thus, we replaced the phenyl ring in NASA with a methylene group to reduce its size without perturbing lysine reactivity (Fig. 5D,F). These and similar analyses led us^104^ to develop a series of ∼30 group-transfer handles (K1-K30) with tunable reactivity (Fig. 5G) by systematically varying the nature of the *N*-alkylation substituents. Similar handles and reactivity trends were reported independently by Hamachi and co-workers.^19^

To develop lysine-based GRIPs, we focused on *N*-<u>su</u>lfonyl-*N*-trifluoromethyl ethan<u>a</u>mide (SuFA) handle with ─CF_3_ because of its minimalist nature, synthetic convenience, and balanced reactivity/stability profile. We generated BRD4 GRIPs from the kinase inhibitors Bosutinib and Ibrutinib scaffolds, targeting EphA2 and BTK, respectively (Fig. 5H,I).^105, 106^ We performed LC-MS/MS analysis to confirm that the alkyne probe **5.3** selectively labeled only Lys617 on EphA2, while compound **5.5** labeled Lys406 on BTK (Fig. 5J,K, and Fig. MS5,6). We synthesized the corresponding BRD4-targeting GRIPs (**5.4** and **5.6**, Fig. 5H,I) and confirmed that the binders exhibit low ATP-pocket occupancy in the BRET assay (Fig. 5L). Finally, we confirmed GRIPs mediated BRD4 phosphorylation in cells transiently expressing BRD4 and EphA2 or BTK, and this phosphorylation was competed away by excess (*S*)-JQ1 (Fig. 5M,N). Overall, these studies provide a suite of tunable group-transfer handles for lysine and first examples of lysine-based GRIPs for inducing POI phosphorylation.

### GRIPs recruit endogenous AKT to FKBP-Liprin to trigger condensate formation with dose and temporal control

As done with eraser GRIPs, we sought to apply these writer GRIPs to control biological processes. Phosphorylation triggers biomolecular condensate formation^107^ and in the case of Liprin-α3, phosphorylation of serine residues within regions flanked by the Liprin homology (LH2) and Sterile alpha motif (SAM) domains facilitate phase separation that regulates synaptic vesicle exocytosis.^108^

We generated a HEK293T cell line stably expressing mVenus–Liprin-α3 fused to FKBP (Fig. 6A)^7^ and synthesized compounds composed of an AKT binder and FKBP binder connected *via* linkers of varying length and chemotype (Fig. 6B and Extended Data Fig. 5A). Automated image-based screening in mVenus–Liprin-α3–FKBP expressing HEK293T cells revealed that GRIP **6.1** induced robust and dose-dependent droplet formation.

**Figure 6.**
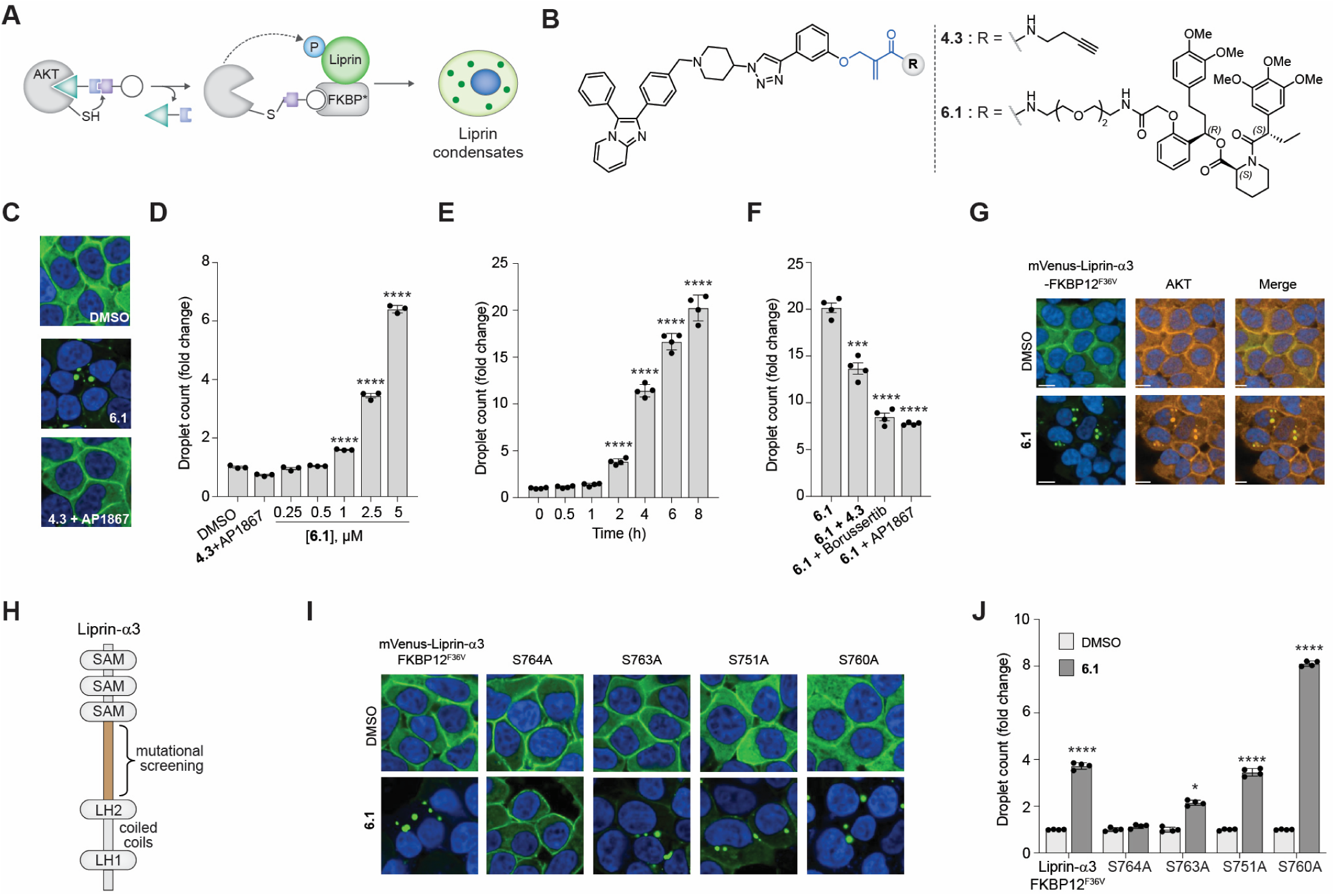
GRIPs can induce phase separation of Liprin. (**A**) Schematic of GRIPs-induced Liprin phosphorylation. (**B**) Structures of Liprin targeting GRIP **6.1** recruiting AKT and corresponding alkyne probe **4.3**. (**C**) Confocal microscopy images (63× objective) of HEK293T cells stably expressing FKBP-tagged Liprin, treated with DMSO, **6.1** (5 μM), or a combination (1:1 ratio) of alkyne probe **4.3** and FKBP binder *o*-AP1867 (5 μM). (**D, E**) Dose- (**D**) and time-dependent (**E**) induction of phase-separated droplets by GRIP **6.1**; unpaired *t*-test, two-tailed. For panel (**E**) 5 μM of **6.1** was used. (**F**) Competition of GRIP **6.1** (2.5 μM) induced phase separation by alkyne probe **4.3**, AKT inhibitor Borussertib, or *o*-AP1867 (5 μM); unpaired *t*-test, two-tailed. (**G**) Microscopic images (40× objective) showing the colocalization in Liprin and AKT in phase-separated droplets. (**H**) Schematic representation of Liprin-α3 with disorder region between coiled-coil regions and sterile alpha motifs (SAM) scanned for phospho-resistant mutational analysis. (**I, J**) Confocal images (**I**) and quantification (**J**) of GRIP **6.1** (5 μM)-induced Liprin-α3 droplets in HEK293T stable cell lines expressing single alanine variants of mVenus–Liprin-α3–FKBP; unpaired *t*-test, two-tailed. Data in panels **D–F**, and **J** represent mean ± SD, (n = 3 for **D**; n = 4 for **E, F,** and **J**).

In contrast, treatment with individual binders or their equimolar combinations failed to induce droplet formation (Fig. 6C,D, and Extended Data Fig. 5B). Phase separation was rapid, with droplets appearing within 2 hours post-treatment (Fig. 6E), enabling temporal control over condensate formation. To confirm the GRIPs mechanism, we performed competition experiments, co-treating **6.1** with AKT inhibitors (**4.3** or Borussertib) or FKBP competitor and observed a decrease in droplet formation (Fig. 6F). Using confocal microscopy, we further demonstrated the colocalization of endogenous AKT with GRIP **6.1**–induced Liprin-α3 condensates, confirming kinase recruitment to the condensate (Fig. 6G). To verify that the observed phenotypic effect arose from action of GRIPs on Liprin-α3, we generated a panel of Liprin-α3 alanine variants (S/T→A) targeting residues between the LH2 and SAM domains, guided by the reported Liprin-α3 phosphosites that trigger droplet formation (Fig. 6H).^7, 108^ Among the alanine variants tested, S764A showed the strongest resistance to GRIP **6.1**–mediated droplet formation (Fig. 6I,J). Interestingly, S760A or S763A variants alone did not abolish condensates, but composite mutations incorporating S764A (e.g., S760A/S764A or S763A/S764A) fully suppressed GRIP activity (Extended Data Fig. 5C,D).

Concurrent with this effort, we developed a phosphorylation-inducing chimera formed by connecting the FKBP binder to the activator of AMP-activated protein kinase (AMPK, Extended Data Fig. 5E). This chimera also induced condensate formation of FKBP-Liprin,^7^ allowing a comparison of GRIPs and conventional chimeras in an identical system. Both GRIPs and AMPK chimera enabled dose- and temporal-controlled condensate formation with similar potency, and their effects were competed out by excess kinase or FKBP binders. Both chimeras did not perturb the global protein expression (determined by a global quantitative proteomics experiment; Extended Data Fig. 5F-H), but their effects on the global phosphoproteome were markedly different. While the global phosphoproteome was mostly unaltered by AKT GRIPs (Extended Data Fig. 5I), the AMPK-chimera dramatically elevated levels of several phosphoproteins (90 phosphopeptides detected, p < 0.01, fold-change >2, Extended Data Fig. 5J), owing to AMPK activation (Extended Data Fig. 5K). Elevated levels of several Liprin phosphosites were observed for the AMPK chimera, validating the proteomics workflow. However, for both the AKT and AMPK chimera, the sites implicated by alanine-scanning studies were undetectable in global phosphoproteomic studies. These sites are surrounded by lysine or arginine and may be furnishing short tryptic peptides that are often undetected in a global phosphoproteomics experiment. Alternatively, condensate-based phosphorylated Liprin may not be accessible. Overall, the global proteomics experiments confirm that, while a conventional chimera with an AMPK activator perturbs the global proteome, the AKT GRIPs do not. Furthermore, these studies report the first examples of dose- and time-controlled Liprin condensate formation using the inhibitor-based GRIPs platform.

### Fully endogenous GRIPs can switch on EGFR signaling

Building on these studies of the hemi-endogenous Liprin system, we set out to develop fully endogenous GRIPs to activate EGFR signaling. Upon EGF binding, EGFR oligomerizes and autophosphorylates tyrosine residues that serve as docking sites for signaling proteins that regulate cellular growth and proliferation.^109^ We envisioned GRIPs that dimerize EGFR to induce autophosphorylation. FKBP-bearing receptor tyrosine kinases and a dimer of FKBP binders have been used extensively to oligomerize and switch on receptor tyrosine kinase signaling ^110, 111^ We envisioned that GRIPs could accomplish similar receptor oligomerization and activation, but on endogenous receptor tyrosine kinases (Fig. 7A). To achieve this, we synthesized potential GRIPs from the EGFR inhibitor Osimertinib^112^ (Fig. 7B, Extended Data Fig. 6A,B). EGFR GRIPs **7.1** induced autophosphorylation in A431 cells at the key Y845 site on the activation loop as well as at additional sites (Y992, Y1045, Y1086, and Y1148) that serve as docking sites for downstream effectors (Fig. 7C). These autophosphorylations were accompanied by endolysosomal degradation of EGFR, as has been observed previously^113, 114^ and an increase in the phosphorylation levels of downstream effectors, including AKT, MEK, and ERK (Fig. 7D). Treatment with **7.1** in cells expressing the kinase-dead EGFR (K745M variant)^115^ did not result in auto-phosphorylation, confirming the on-target selectivity (Extended Data Fig. 6C,D).

**Figure 7.**
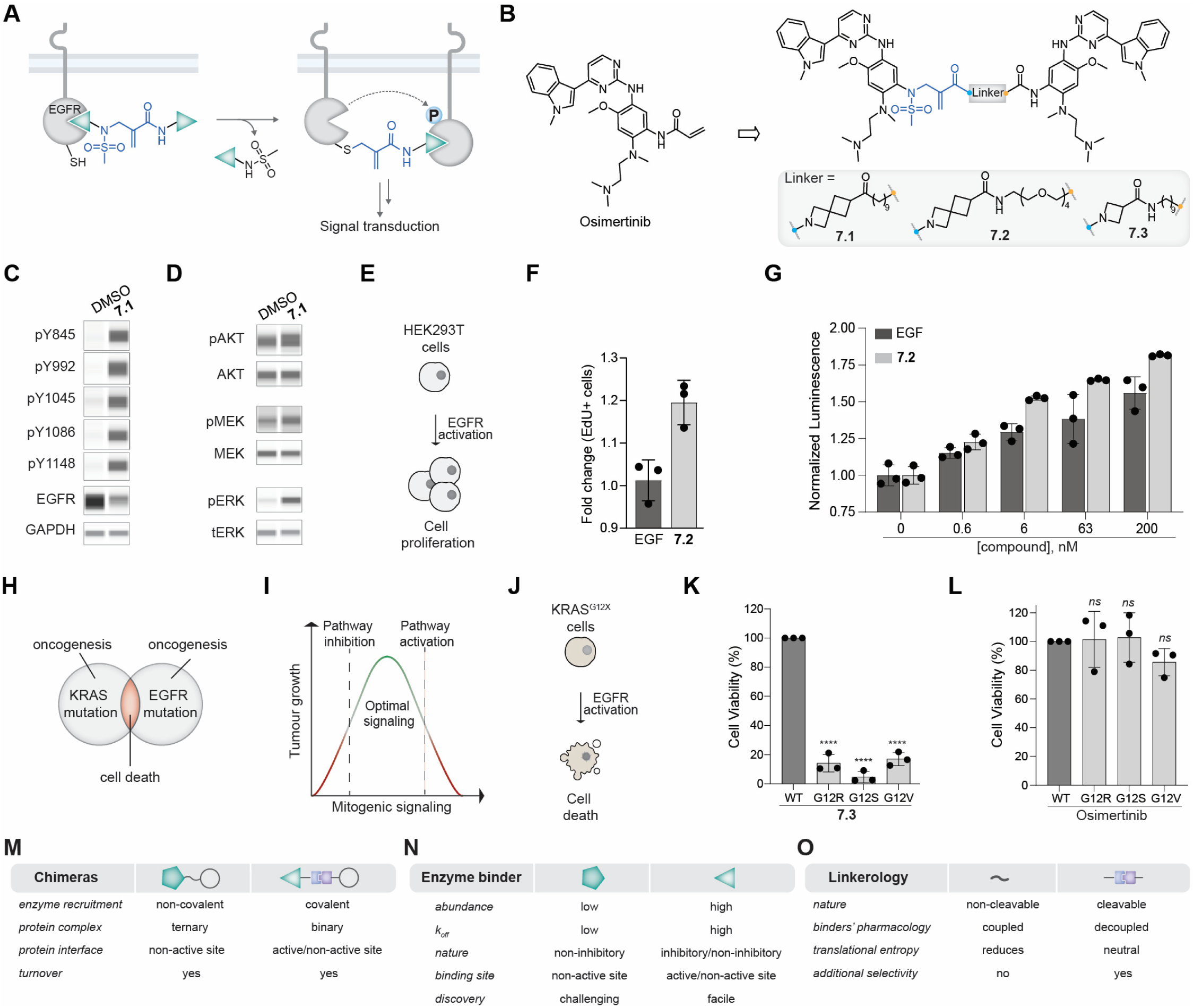
GRIPs switch EGFR signaling on. (**A**) Schematic of EGFR pathway activation by GRIPs. (**B**) Structure of EGFR inhibitor Osimertinib utilized to generate EGFR GRIPs **7.1–3.** (**C,D**) EGFR phosphorylation (**C**) by GRIP **7.1** (50 nM) and downstream signaling activation (**D**). (**E**) Schematic of the induction of HEK293T cell proliferation by GRIP **7.2** or EGF. (**F**) Fold change of Edu+ HEK293T cells treated with GRIP **7.2** or EGF (250 nM) measured via confocal imaging. (**G**) Quantitation of nLuc-CRBN expression in HEK293T. For panels **F-G**, GRIP treatments (n = 3) are normalized to the mean value of DMSO; unpaired t-test, two-tailed. Bars represent mean ± SD. (**H**) Schematic showing synthetic lethality of KRAS and EGFR mutations. (**I,J**) Rationale of TOVER (**I**) and its application in KRAS mutant cells (**J**). (**K,L**) Cell viability of KRAS G12R (PSN1), G12S (A549), and G12V (NCIH727) compared to KRAS WT (NCIH1299) cell lines with compound **7.3** (**K**) or Osimertinib (**L**) treatment (7 μM, 2% v/v DMSO, 72 h). Viability was determined by normalization to DMSO only of KRAS WT line. Bars represent mean ± SD (n = 3). Statistical significance compared to KRAS WT was determined by an ordinary one-way ANOVA test (**** p < 0.0001 and ns p >0.05). (**M–O**) Key differences between the traditional design and GRIPs on the chimera (**M**), enzyme binder (**N**), and linker levels (**O**).

We next sought to use EGFR-activating GRIPs to stimulate cell proliferation and thus protein production by mimicking EGF signaling (Fig. 7E). To test this, we treated HEK293T cells, commonly used for protein production, with GRIP **7.2** or EGF (250 nM) for 12 h in serum-free media. In agreement with reported FKBP-engineered receptors,^110^ treatment with GRIP **7.2** resulted in a ∼20% increase in cell proliferation compared with EGF treatment, as measured by EdU incorporation using confocal microscopy (Fig. 7F and Extended Data Fig. 6E).^116^ This was further validated using HEK293T cells expressing luciferase (nLuc-CRBN) as a reporter system for protein expression. Here, we observed a dose-dependent increase (up to ∼75%) in nLuc-CRBN in cells treated with **7.2**, consistent with the trend observed with EGF treatment (Fig. 7G). Similar results were obtained in engineered CHO cells using an orthogonal flow cytometry readout (Extended Data Fig. 6F-H).

Using EGFR GRIPs, we also pharmacologically validate the genetic observations that EGFR-activating mutations are synthetically lethal to cells with oncogenic KRAS mutations (Fig. 7H). Such synthetic lethality potentially arises because cancer cells have Goldilocks levels of signaling, and therapeutic overactivation of oncogenic signaling (TOVER) is detrimental to cancer cells (Fig. 7I,J).^43, 117^ Using the EGFR switch-on GRIP **7.3**, we observed selective lethality in KRAS mutant cells but not in wild-type cells (Fig. 7K). In contrast, the EGFR covalent inhibitor Osimertinib showed no such effect (Fig. 7L), providing preliminary evidence that GRIPs can achieve therapeutic overactivation of oncogenic signaling.^43^ Collectively, these studies report the first endogenous approach to activate EGFR signaling via dimerization and subsequent phosphorylation. These GRIPs can serve as proteolytically stable mimics of EGF in biomanufacturing and tools to study TOVER in oncology.

## Discussion

We report a scalable platform for modulating PTMs by repurposing inhibitors of writers/erasers. In contrast to conventional chimeras that rely on scarce, non-inhibitory binders and often activators, GRIPs utilize abundant, high-quality inhibitors—their abundance enables platform scalability, while their high-quality increases GRIPs’ translational potential. Furthermore, we provide a toolbox of ∼42 group-transfer handles and > 5000 compatible inhibitor scaffolds. GRIPs differ from conventional chimeras in several ways, including the nature of linkers and mechanisms (Fig. 7M-O). As conventional chimeras lack cleavable handles (Fig. 1A), they do not uncouple the pharmacology of the two binders. In contrast, GRIPs allow the release, or “washing out,” of one of the binders and its pharmacological effects. Furthermore, GRIPs’ covalent recruitment of writer/eraser may improve turnover, as we recently reported using chemogenetic tags^83^, and may reduce their susceptibility to efflux pumps, an issue with several chimeras.^118^ GRIPs still require high-quality inhibitors that are currently unavailable for several writers and erasers, and the *in vivo* efficacy and safety of GRIPs is yet to be determined.

The GRIPs platform provides a multifaceted approach to modulate PTMs. They can be used as molecular switches to turn on/off signaling. We anticipate *that O-GlcNAc GRIPs will open new avenues of biological interrogation for this nascent field and will also* propel the development of GRIPs for other PTMs, leveraging the ready availability of high-quality inhibitors. Beyond these tools, GRIPs endowed new activities to several known drugs, including removing rebound signaling and improving potency and persistence over occupancy-driven drugs. Finally, EGFR GRIPs mimic growth factors such as EGF, offering a proteolytically stable and cost-effective alternative to unstable, expensive growth factors in biomanufacturing applications.^119–121^ Overall, the GRIPs platform’s modularity, scalability and reliance on existing inhibitors position it well for probing biological systems and propelling the development of novel therapeutic modalities.

## Data availability

The PDB mining methods are available at https://github.com/as1000/LigandableResiDist. Additional supporting information generated during the current study but not included in the manuscript are available from the corresponding author upon request.

## Acknowledgments

This work was supported by the Merkin Institute of Transformative Technologies in Healthcare, DARPA (HR0011-21-2-0010), NIH (R21AI154099, R21AI178690, U01DK137242, R01GM137606) (to A.C), NIH K99GM159057 (to E.K), the Prince Alwaleed Bin Talal Research Fellowship from the Dubai Harvard Foundation for Medical Research (to S.H.S), the Damon Runyon Cancer Research Foundation, Suzanne and Bob Wright Fellowship DRG-2539-24 (to C.L), the Natural Sciences and Engineering Research Council of Canada 587836-2024 (to K.K), NIH R35GM138014 (to N.H.S.), National Science Foundation CHE-2235508 (to C.F), Arnold and Mabel Beckman Foundation Young Investigator Award (to K.B.), and Packard Foundation Fellowship (to K.B.).

## Author contributions

E.K. V.V, S.S, J.S, and C.L contributed equally to this work and will put their name first in their curricula vitae citation or elsewhere. R.P, V.S, and K.K contributed equally to this. They will put their name first in their curriculum vitae citations or elsewhere.

## Competing financial interests

Broad Institute has filed patent applications for the work described herein, some of which were licensed to Photys Therapeutics. A.C. is the scientific founder and is on the scientific advisory board of Photys Therapeutics. J.K. is an employee of Daiichi Sankyo Co., Ltd.

## Extended Data Figures

**Extended Data Figure 1.**
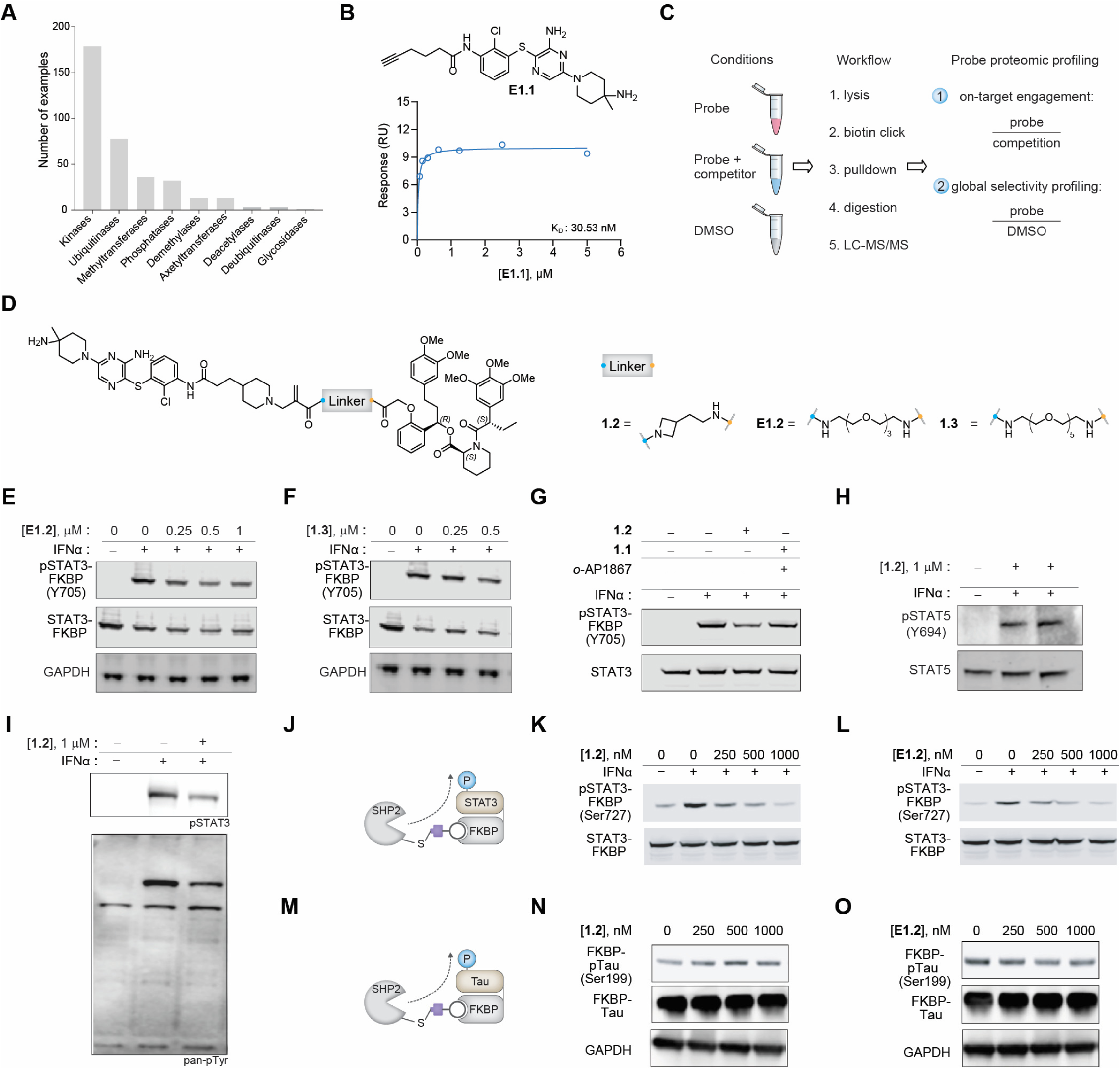
Validation of the SHP2 GRIPs platform. (**A**) Distribution of cysteine targeting covalent fragment chemotypes identified from global proteomic profiling among various classes of writers/erasers. (**B**) Structure of SHP099 modified binder **E1.1** and SPR binding analysis of **E1.1** to recombinant SHP2. (**C**) Global proteomics workflow to determine GRIPs on-target engagement and global selectivity profiling. (**D**) Structures of SHP2 recruiting GRIPs (**1.2**, **1.3,** and **E1.2**). (**E,F**) Dephosphorylation of IFNα-induced STAT3 phosphorylation with GRIPs **E1.2** (**E**), and **1.3** (**F**). (**G**) STAT3 dephosphorylation by GRIPs **1.2** (1 μM) and the equimolar mixture (1 μM each) of SHP2 alkyne probe **2.1** and *o*-AP1867. (**H**) Western blot analysis of endogenous pSTAT5 upon induction with IFNα and treatment with **1.2**. (**I**) Western blot analysis of pSTAT3 and pan-pTyr upon induction with IFNα and treatment with **1.2**. (**J**-**L**) Schematic of STAT3-FKBP ser/thr dephosphorylation (**J**) and western blot analysis upon treatment with GRIPs **1.2** (**K**) and **E1.2** (**L**). In all relevant panels, STAT3/5 phosphorylation is induced by 100 ng/mL IFNα. (**M**-**O**) Schematic of dox-inducible FKBP-tau ser/thr dephosphorylation (**M**) and western blot analysis upon treatment with GRIPs **1.2** (**N**) and **E1.2** (**O**)

**Extended Data Figure 2.**
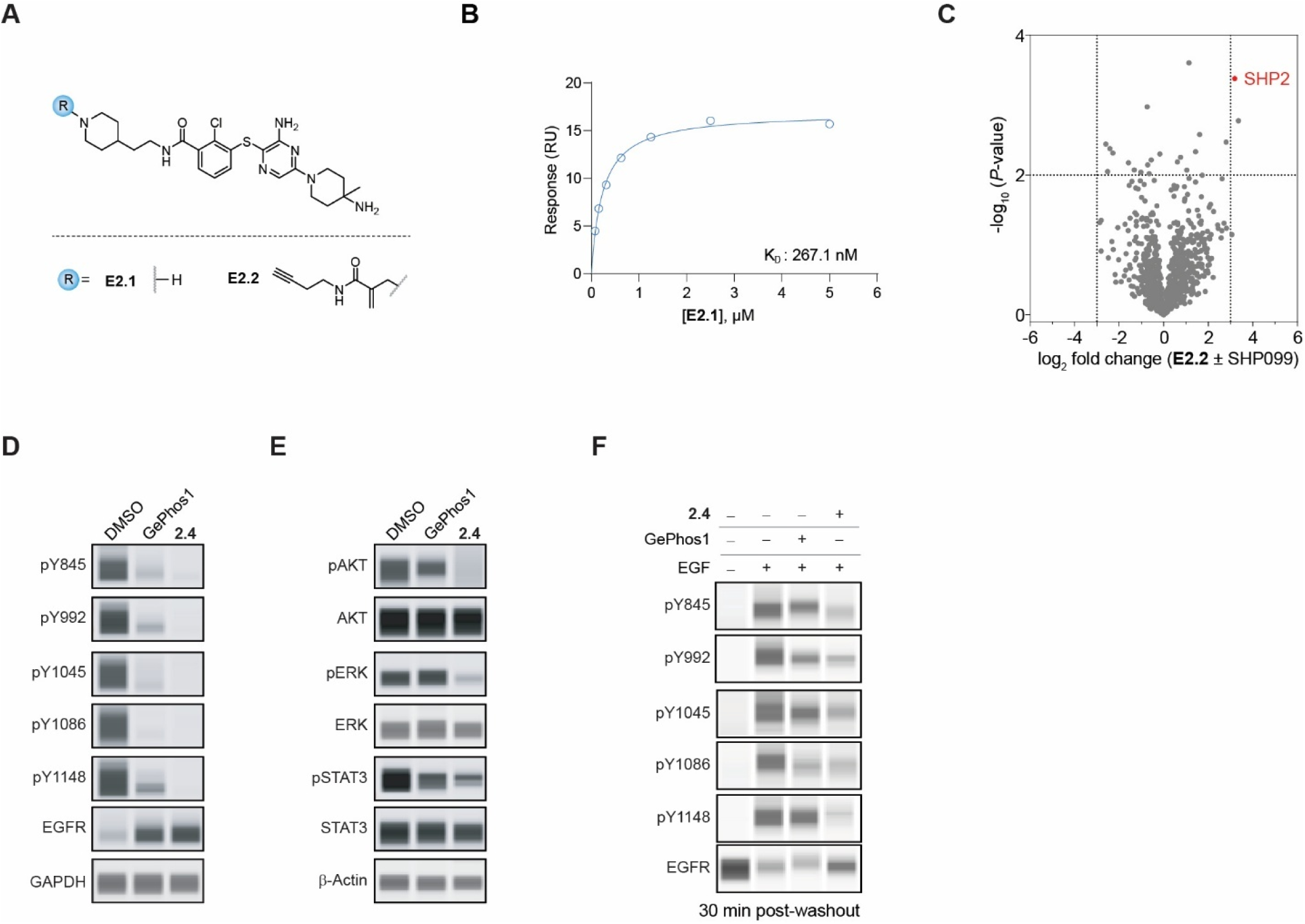
Endogenous applications of the SHP2 GRIPs platform. (**A**) Structures of modified SHP2 scaffold (**E2.1**) and alkyne SHP2 GRIP (**E2.2**) with reverse amide modification. (**B**) SPR binding analysis of **E2.1** to recombinant SHP2. (**C**) Global proteomic profiling of labeling with probe **E2.2** (100 nM) in competition with SHP099 (500 nM). Thresholds are defined at fold change [**E2.2**/competition] = 8 and p value < 0.01. (**D,E**) EGFR dephosphorylation (**D**) and downstream signaling inhibition (**E**) of HeLa cells treated with control inhibitor GePhos1 and **2.4** (6h treatment of 250 nM each). (**F**) Western-blot assay to monitor pEGFR and EGFR levels of HeLa cells treated with **2.4** or GePhos1 (treatment of 250 nM each) 30 min post-washout.

**Extended Data Figure 3.**
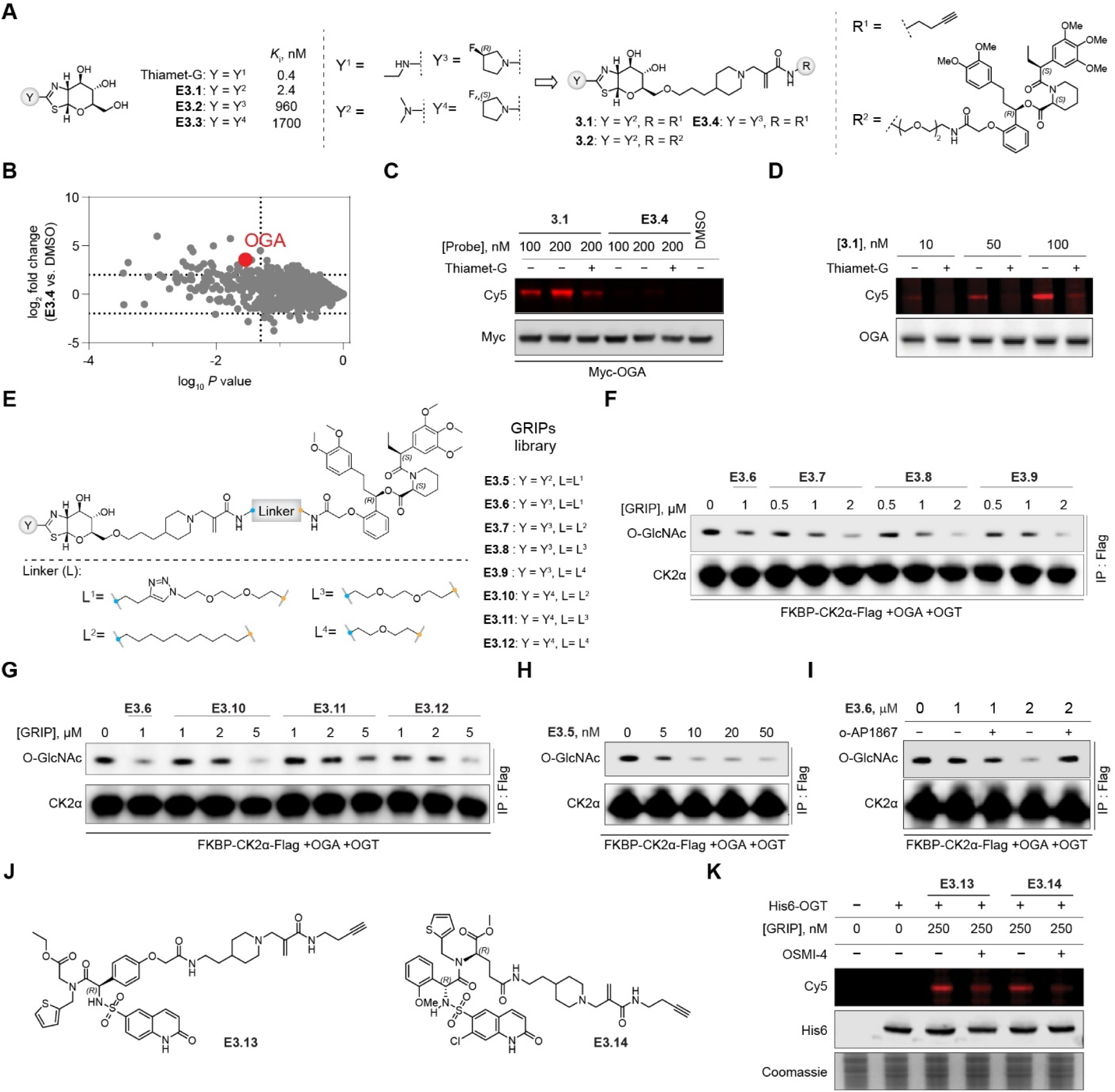
Validation of OGA and OGT GRIPs. (**A**) Structure of Thiamet-G and analogs with varying Ki values and structures of alkyne (**3.1** and **E3.4**) or o-AP1867 (**3.2**)-based GRIPs. (**B**) Global proteomic profiling of alkyne probe **E3.4**. Thresholds are defined at fold change [probe/control] = 4 and p value < 0.05. (**C**) Cellular labeling of overexpressed OGA by GRIPs **3.1** and **E3.4** and competition with Thiamet-G (2 μM). (**D**) Dose and competition-dependent labeling of endogenous OGA by GRIP **3.1** and competition with Thiamet-G (500 nM) (**E**) Library of o-AP1867-based GRIPs with varying Thiamet-G analogs and linkers (**E3.5-12**). (**F-G**) Dose-dependent de-*O*-GlcNAcylation of FKBP-CK2α using GRIPs **E3.6-9** (**F**) and **E3.6-E3.10-12** (**G**). (**H**) Dose-dependent de-*O*-GlcNAcylation of FKBP-CK2α using GRIPs **E3.5**. (**I**) Dose- and competition-dependent de-*O*-GlcNAcylation of FKBP-CK2α using GRIPs **E3.6**. (**J**) Structures of OGT alkyne GRIPs **E3.13-14**. (**K**) Labeling of His6-OGT by alkyne GRIPs **E3.13-4**.

**Extended Data Figure 4.**
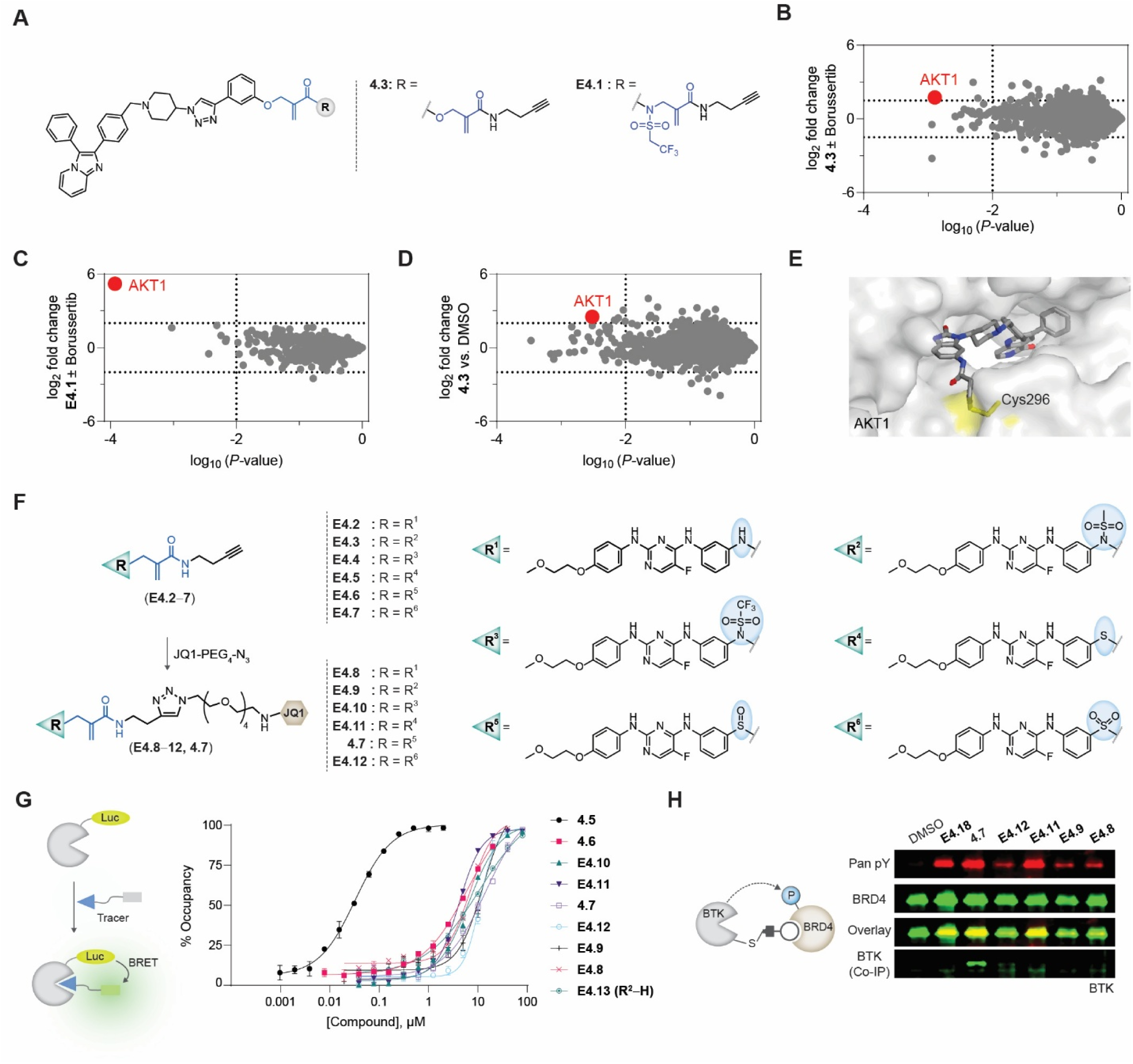
Validation of the GRIPs phosphorylation platform. (**A**) Structures of AKT alkyne GRIP probes (**4.3** and **E4.1**) utilized for target engagement studies from an allosteric covalent AKT inhibitor. (**B,C**) Competition-based target engagement studies using global proteomic profiling for **4.3** (**B**) and **E4.1** (**C**). (**D**) Selectivity profiling of probe **4.3** using global proteomic profiling. DMSO-treated cells are used for comparison. For panels B-D thresholds are defined as fold change [probe/control] > 4, p value < 0.01 (**E**) Crystal structure of AKT1 in Complex with Covalent-Allosteric AKT Inhibitor Borussertib (PDB ID: 6HHF). Targeted cysteine (C296 is highlighted in yellow). The other targeted cysteine (C310) was not resolved in this crystal structure. (**F**) Synthetic scheme of Sperbutinib alkynes (**E4.2-7**) converted to the corresponding GRIPs (**E4.8-12** and **4.7,** *left*). Structures of alkyne analogs and corresponding GRIPs for Sperbutinib-based binders and GRIPs with derivatized or replaced aniline groups (*right*). (**G**) Nano-BRET assay for Sperbutinib-based binders and GRIPs with a derivatized or replaced aniline group, as well as covalent (**4.5**), non-covalent (**4.6**), and non-covalent sulfonamide (**E4.12**) Sperbutinib compounds. Data are presented as mean values with standard errors; n = 2. (**H**) BRD4 phosphorylation assay for BTK recruiting GRIPs **E4**.**8-12** and **4.7**. For panel I, 250 nM of GRIPs is used and 1 μM of (S)-JQ1 for competition.

**Extended Data Figure 5.**
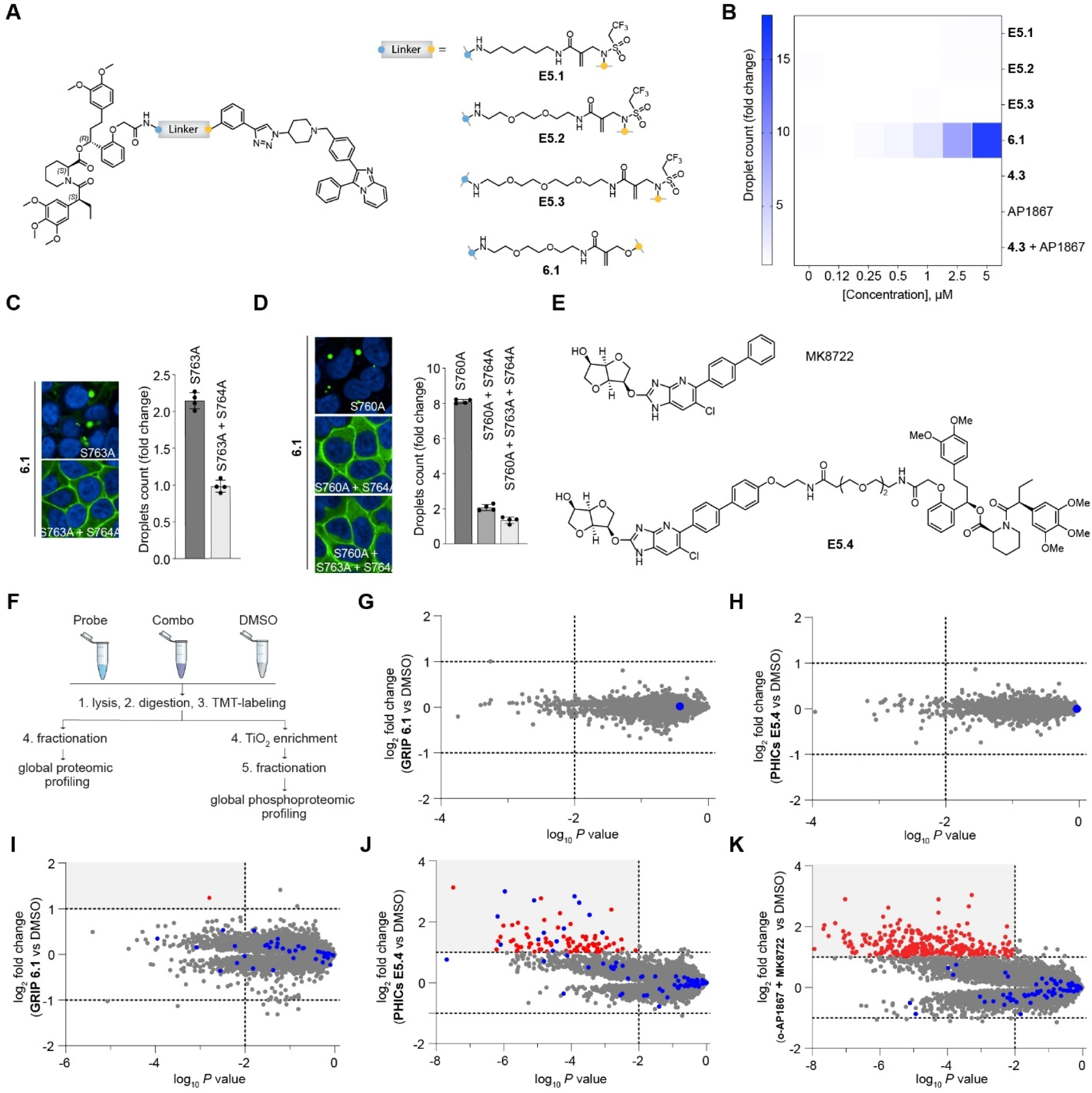
Optimization and validation of AKT-GRIPs for the phase-separation of Liprin. (**A**) Structures of GRIPs **6.1** and **E5.1-3**. (**B**) Dose-dependent induction of Liprin droplets by GRIPs (**6.1** and **E5.1-3)**, monomeric components (**4.3** and *o*-AP1867), and mixture (1:1 of **4.3** and *o*-AP1867). Data are presented as a heat map of the mean value of n = 4. (**C-D**) Mutational analysis of S763A in combination with S764A (**C**) and S760A in combination with S764A and S764A+S763A (**D**) after treatment with **6.1** (5 μM). Data are presented as mean values ± SD, n = 4. (**E**) Structures of AKT activator MK8722 and PHICs compound **E5.4**. (**F**) Workflow for total proteome (*left*) and phosphoproteome (*right*) for GRIPs and PHICs molecules. (**G,H**) Global proteomic profiling of proteins in FKBP-Liprin-expressing cells with 5 μM GRIP **6.1** (**G**) or 5 μM AMPK-PHICs **E5.4** (**H**) versus DMSO. FKBP-Liprin is colored blue. (**I**-**K**) Global phosphoproteomic profiling showing detected phosphopeptides in FKBP-Liprin-expressing cells treated with 5 μM GRIP **6.1** (**I**), or 5 μM AMPK-chimera PHICs (**J**), or a mixture of PHICs monomeric components (1:1 ratio) (**K**) versus DMSO. FKBP-Liprin phosphopeptides are colored blue, while off-targets are colored red. For panels G-K Thresholds are defined as fold change [treatment/DMSO] > 2, p value < 0.01

**Extended Data Figure 6.**
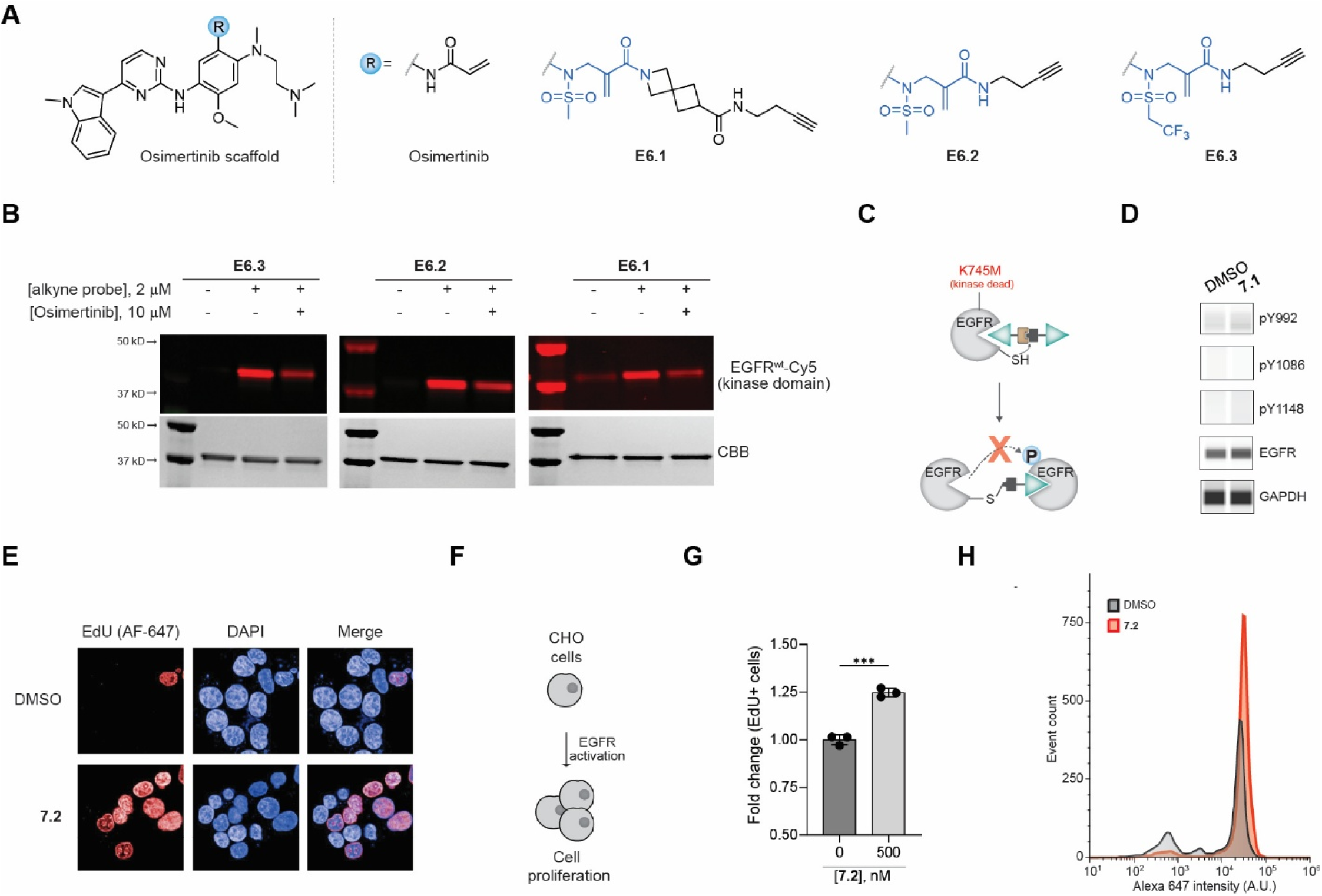
GRIPs switch on/off signaling pathways. (**A**) Structures of Osimertinib-based GRIP alkyne probes **E6.1–3**. (**B**) Labeling of wild-type EGFR kinase domain (EGFR^wt^) with different Osimertinib alkyne probes (**E6.1–3**). Compounds were treated at 2 μM for 2 h at room temperature. (**C**) Schematic of kinase dead (K745M) dimerization by GRIPs. (**D**) EGFR (K745M) phosphorylation levels after treatment with **7.1** (250 nM). (**E**) Representative confocal images of HEK293T cells treated with DMSO or GRIP **7.2** (250 nM). (**F**) Schematic of CHO cell proliferation induced by GRIPs. (**G**) Quantitation of Edu+ ExpiCHO cells treated with GRIP **7.2** (500 nM). Data represent mean ± SD, (n = 3). (**H**) Flow cytometry histograms of ExpiCHO cells treated with DMSO and GRIP **7.2** (500 nM).

