## Supplementary Figures for "A Scalable Design for Proximity-Inducing Molecules"

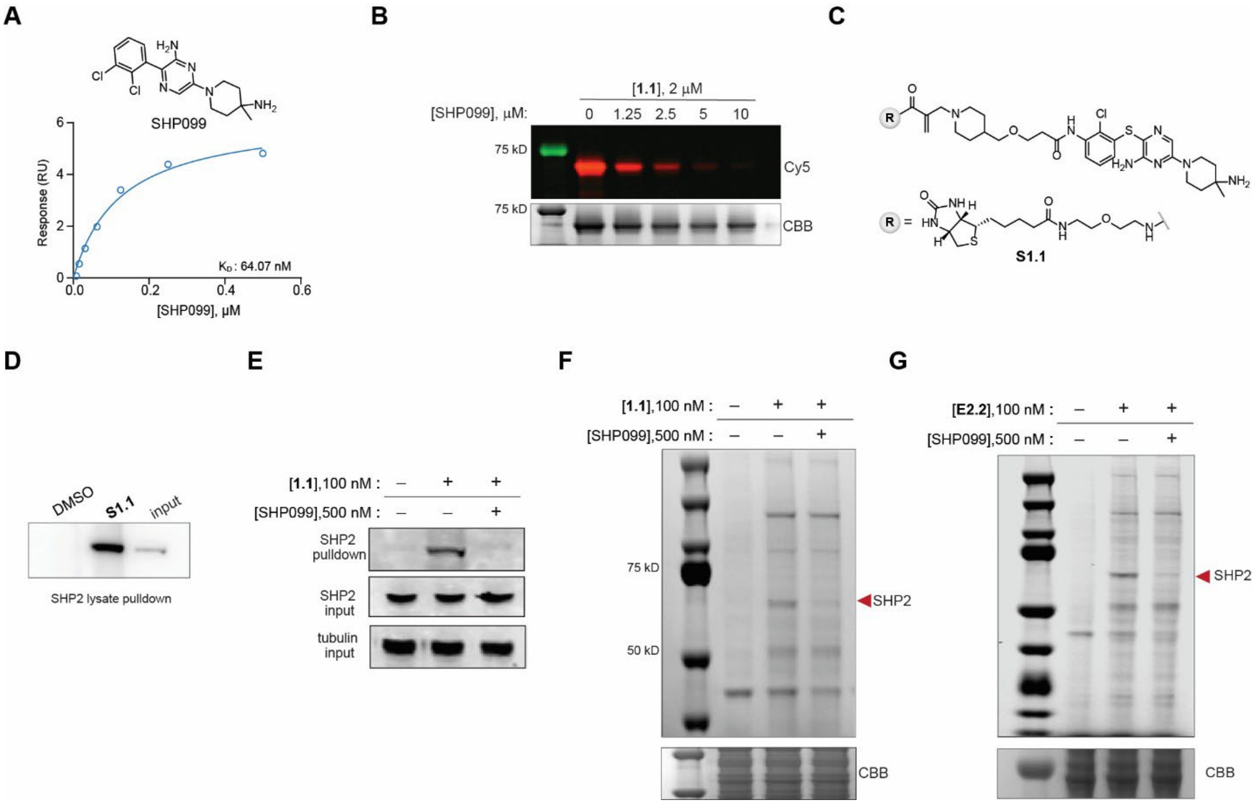


**Supporting Figure 1.** (**A**) SPR binding analysis of SHP099 on recombinant SHP2 protein. (**B**) Biochemical labeling of purified SHP2 using **1.1** and competition with SHP099. CBB = Coomassie brilliant blue staining. (**C**) Structure of biotinylated probe **S1.1**. (**D**) Pulldown of endogenous SHP2 from HEK293T lysate using biotinylated probe **S1.1** (2 μM). (**E**) Pulldown of endogenous SHP2 after cell treatment with alkyne probe **1.1**, followed by cell lysis, copper-click with desthiobiotin-azide, streptavidin-mediated pulldown, and release with excess biotin. (**F**) Cellular labeling using alkyne probe **1.1** and competition with SHP099. Labeling is evaluated by conjugating labeled proteins with Cy5-azide via copper-click chemistry, followed by in-gel fluorescence. SHP2 is approximately 68 kDa. (**G**) In-cell labeling of SHP2 by **E2.2** and competition with SHP099.

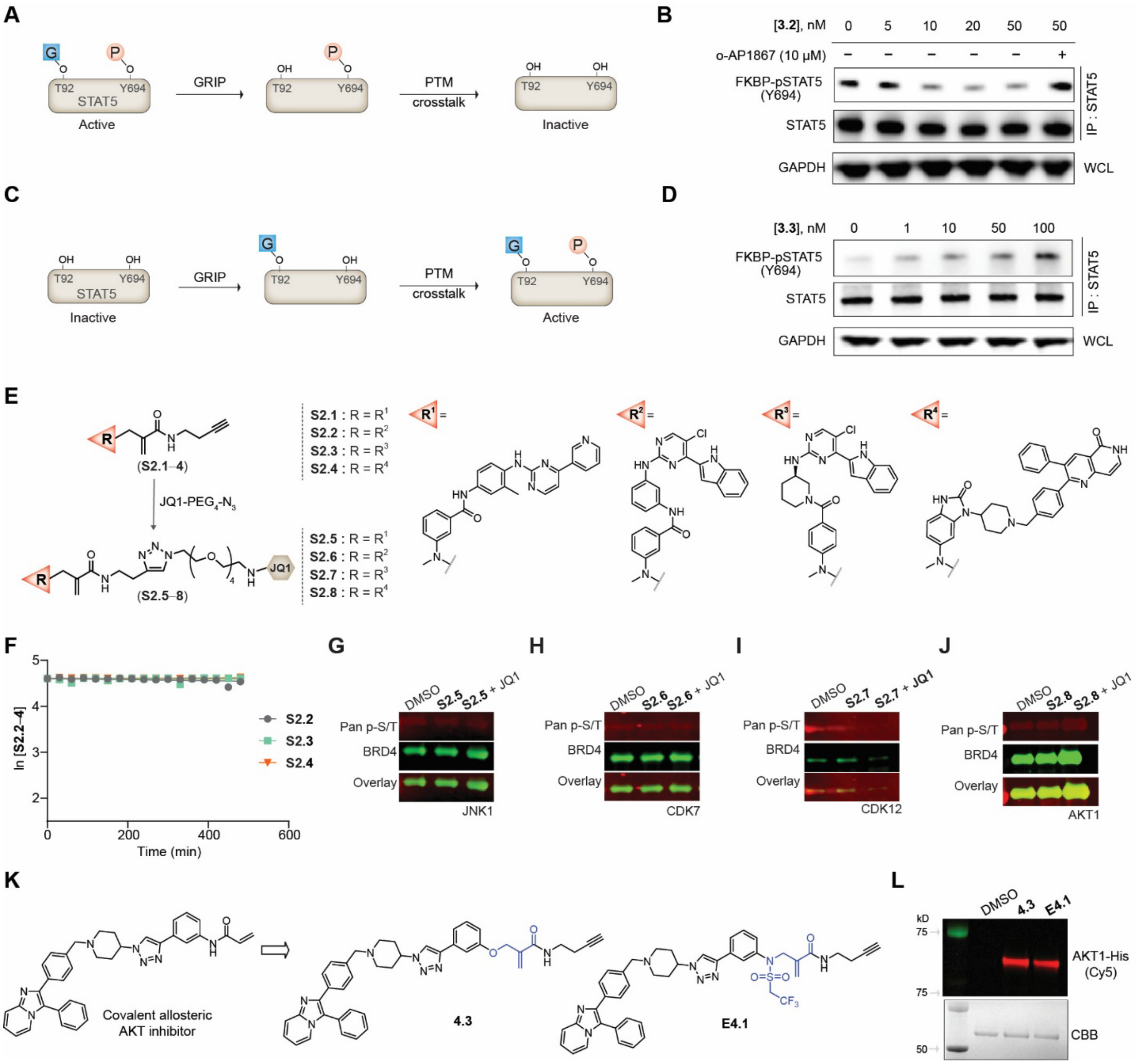


**Supporting Figure 2.** (**A**) Schematic of O-GlcNAc-dependent dephosphorylation of STAT5 Y694 using OGA GRIPs. (**B**) Dose- and competition-dependent dephosphorylation of FKBP$^{F36V}$-STAT5 (pY694) using GRIP **3.2**. (**C**) Schematic of O-GlcNAc-dependent phosphorylation of STAT5 Y694 using GRIPs. (**D**) Dose-dependent phosphorylation of FKBP$^{F36V}$-STAT5 (pY694) using GRIP **3.3**. (**E**) Schematic for converting aniline-based binders bearing alkyne handles (**S2.1-4**) to the corresponding GRIPs (**S2.5-8**) via CuAAC with the corresponding JQ1-azide (left). Structures of aniline-based binders targeting JNK1, CDK7, CDK12, and AKT1 (in order of appearance, right). (**F**) Reaction kinetics of compounds **S2.2-4** with N-Acetyl-Cysteine-OMe ester in PBS. Determination of the corresponding reaction rates was not possible because no reactivity was observed. (**G-J**) BRD4 phosphorylation assay for GRIPs **S2.5-8,** recruiting the corresponding targeting kinase. For all relevant panels 250 nM of GRIPs is used and 1 μM of (S)-JQ1 for competition. (**K**) Generation of AKT alkyne GRIP probes (**4.3** and **E4.1**) utilized for target engagement studies from an allosteric covalent AKT inhibitor. (**L**) Biochemical labeling of purified AKT1 protein (2 μM) using 2 μM of alkyne probes **4.4** and **E4.1**.

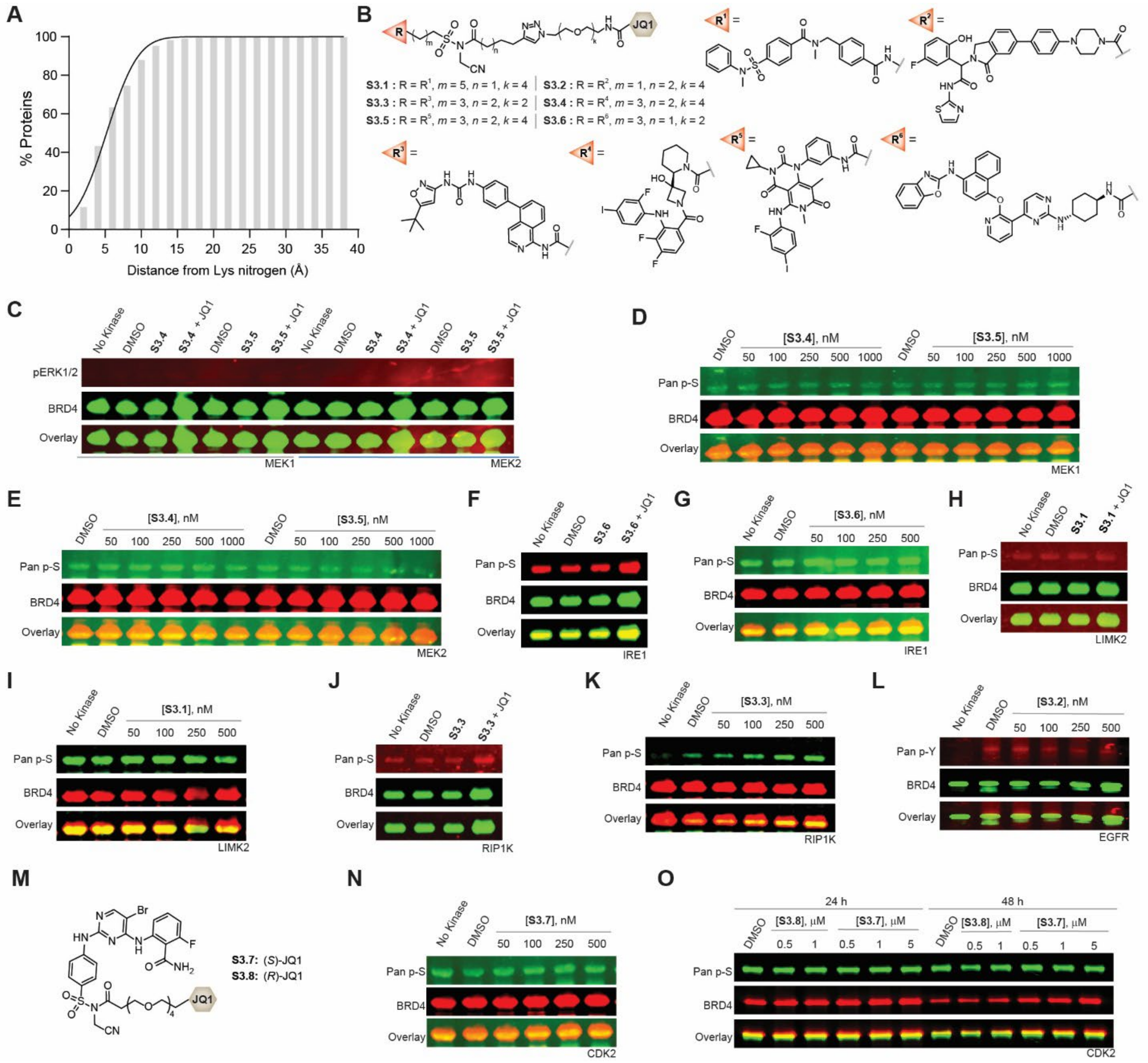


**Supporting Figure 3.** (**A**) Distribution of distances of protein binders from lysines across the human proteome. (**B**) Structures of diverse allosteric kinase inhibitors utilized to generate NASA-based GRIPs **S3.1-6**. (**C–L**) BRD4 phosphorylation assays for GRIPs **S3.1-6** recruiting the corresponding kinases. (**M**) Structures of CDK2 NASA-based GRIPs **S3.7–8** based on the parent ligand for CDK2 kinase. (**N–O**) BRD4 phosphorylation assay for GRIPS **S3.7–8** in a dose and time-dependent manner. For panels **C,F,H,J** 250 nM of GRIPs and for competition 1 μM of (*S*)-JQ1 were used.
