## Supplementary material for "A Scalable Design for Proximity-Inducing Molecules": MS Supplementary Figures

| #1 | b⁺ | b²⁺ | b³⁺ | Seq. | y⁺ | y²⁺ | y³⁺ | #2 |
|---|---|---|---|---|---|---|---|---|
| 1 | 129.06585 | 65.03657 | 43.69347 | Q | | | | 16 |
| 2 | 186.08732 | 93.54730 | 62.70062 | G | 2036.90584 | 1018.95656 | 679.64013 | 15 |
| 3 | 333.15573 | 167.08150 | 111.72343 | F | 1979.88438 | 990.44583 | 660.63298 | 14 |
| 4 | 519.23504 | 260.12116 | 173.74987 | W | 1832.81596 | 916.91162 | 611.61017 | 13 |
| 5 | 648.27764 | 324.64246 | 216.76406 | E | 1646.73665 | 823.87196 | 549.58373 | 12 |
| 6 | 777.32023 | 389.16375 | 259.77826 | E | 1517.69406 | 759.35067 | 506.56954 | 11 |
| 7 | 924.38864 | 462.69796 | 308.80107 | F | 1388.65146 | 694.82937 | 463.55534 | 10 |
| 8 | 1053.43124 | 527.21926 | 351.81526 | E | 1241.58305 | 621.29516 | 414.53253 | 9 |
| 9 | 1154.47892 | 577.74310 | 385.49782 | T | 1112.54046 | 556.77387 | 371.51834 | 8 |
| 10 | 1267.56298 | 634.28513 | 423.19251 | L | 1011.49278 | 506.25003 | 337.83578 | 7 |
| 11 | 1395.62156 | 698.31442 | 465.87870 | Q | 898.40871 | 449.70800 | 300.14109 | 6 |
| 12 | 1523.68013 | 762.34371 | 508.56490 | Q | 770.35014 | 385.67871 | 257.45490 | 5 |
| 13 | 1651.73871 | 826.37299 | 551.25109 | Q | 642.29156 | 321.64942 | 214.76870 | 4 |
| 14 | 1780.78131 | 890.89429 | 594.26529 | E | 514.23298 | 257.62013 | 172.08251 | 3 |
| 15 | 2018.85889 | 1009.93308 | 673.62448 | C-SAS_me... | 385.19039 | 193.09883 | 129.06831 | 2 |
| 16 | | | | K | 147.11280 | 74.06004 | 49.70912 | 1 |

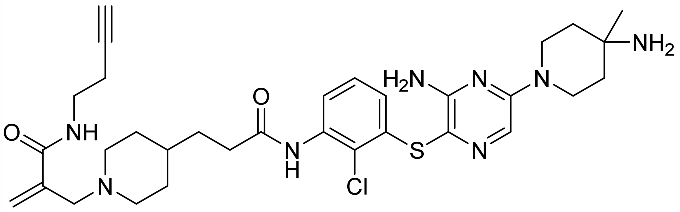

**1.1**

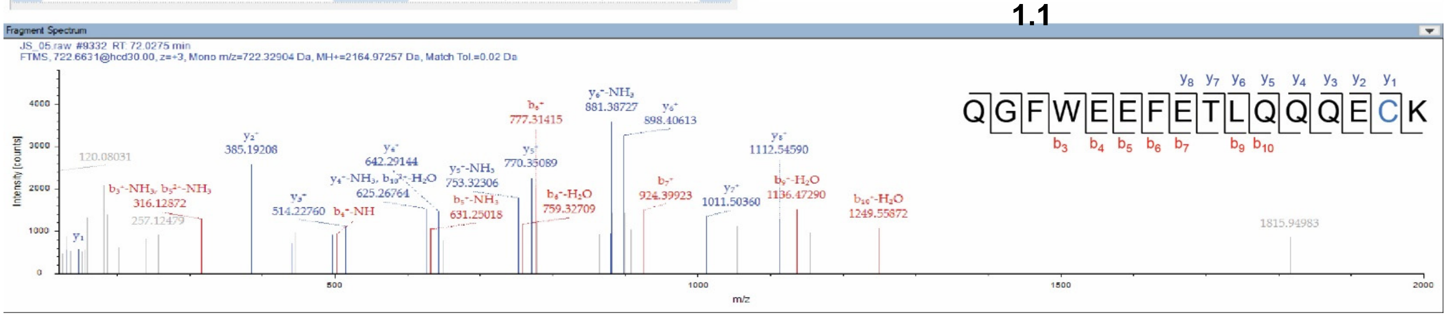


**Figure MS1.** Annotated MS/MS spectrum of a labeled peptide with alkyne probe **1.1** on SHP2**,** including peptide coverage and labeled Cys259.

| #1 | b+ | b2+ | Seq. | y+ | y2+ | #2 |
|---|---|---|---|---|---|---|
| 1 | 114.09134 | 57.54931 | L | | | 17 |
| 2 | 227.17540 | 114.09134 | I | 1965.02623 | 983.01675 | 16 |
| 3 | 324.22817 | 162.61772 | P | 1851.94217 | 926.47472 | 15 |
| 4 | 480.32928 | 240.66828 | R | 1754.88940 | 877.94834 | 14 |
| 5 | 593.41334 | 297.21031 | L | 1598.78829 | 799.89778 | 13 |
| 6 | 721.50831 | 361.25779 | K | 1485.70423 | 743.35575 | 12 |
| 7 | 778.52977 | 389.76852 | G | 1357.60926 | 679.30827 | 11 |
| 8 | 877.59818 | 439.30273 | V | 1300.58780 | 650.79754 | 10 |
| 9 | 990.68225 | 495.84476 | L | 1201.51939 | 601.26333 | 9 |
| 10 | 1091.72993 | 546.36860 | T | 1088.43532 | 544.72130 | 8 |
| 11 | 1205.77285 | 603.39007 | N | 987.38764 | 494.19746 | 7 |
| 12 | 1302.82562 | 651.91645 | P | 873.34472 | 437.17600 | 6 |
| 13 | 1416.86854 | 708.93791 | N | 776.29195 | 388.64961 | 5 |
| 14 | 1654.94613 | 827.97670 | C-SAS_me... | 662.24903 | 331.62815 | 4 |
| 15 | 1783.98872 | 892.49800 | E | 424.17144 | 212.58936 | 3 |
| 16 | 1931.05714 | 966.03221 | F | 295.12885 | 148.06806 | 2 |

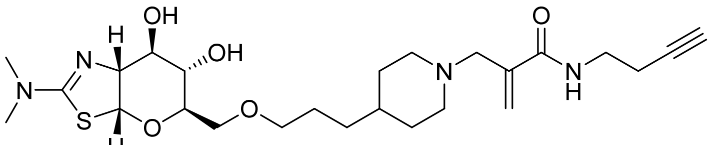

**3.1**

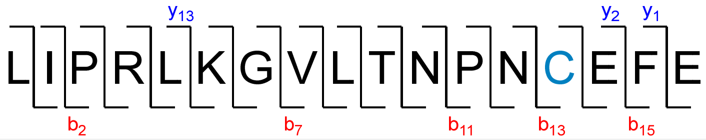


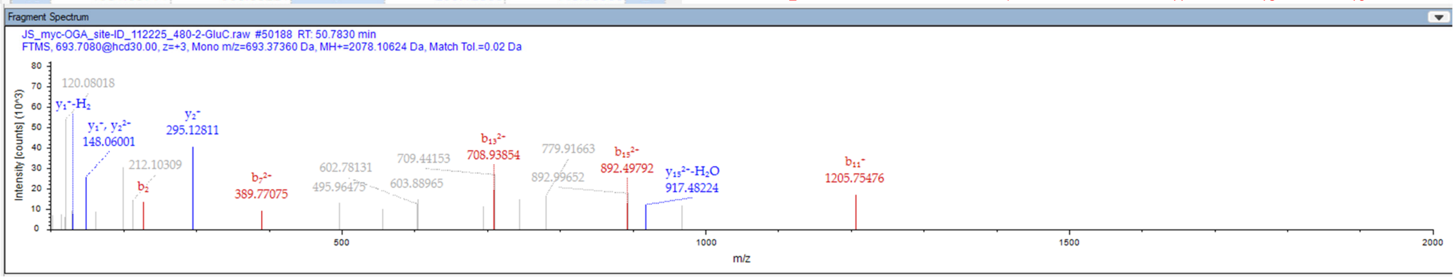


**Figure MS2.** Annotated MS/MS spectrum of a labeled peptide with alkyne probe **3.1** on OGA**,** including peptide coverage and labeled Cys316.

OGT1(397–408)

| #1 | b⁺ | b²⁺ | Seq. | y⁺ | y²⁺ | #2 |
|---|---|---|---|---|---|---|
| 1 | 130.04987 | 65.52857 | E | | | 12 |
| 2 | 261.09035 | 131.04882 | M | 1390.61297 | 695.81012 | 11 |
| 3 | 389.14893 | 195.07810 | Q | 1259.57248 | 630.28988 | 10 |
| 4 | 504.17587 | 252.59158 | D | 1131.51391 | 566.26059 | 9 |
| 5 | 603.24429 | 302.12578 | V | 1016.48696 | 508.74712 | 8 |
| 6 | 731.30287 | 366.15507 | Q | 917.41855 | 459.21291 | 7 |
| 7 | 788.32433 | 394.66580 | G | 789.35997 | 395.18362 | 6 |
| 8 | 859.36144 | 430.18436 | A | 732.33851 | 366.67289 | 5 |
| 9 | 972.44551 | 486.72639 | L | 661.30140 | 331.15434 | 4 |
| 10 | 1100.50408 | 550.75568 | Q | 548.21733 | 274.61230 | 3 |
| 11 | 1338.58167 | 669.79447 | C-SAS_me... | 420.15875 | 210.58302 | 2 |
| 12 | | | Y | 182.08117 | 91.54422 | 1 |

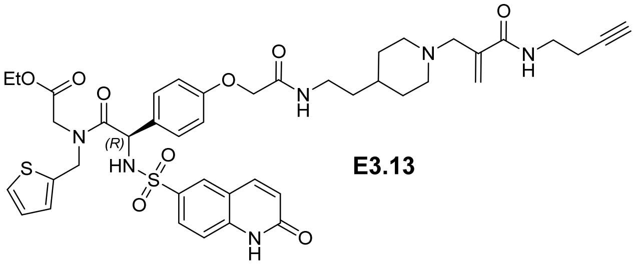

**E3.13**

JS_CL_OGT_site-ID_112225_2240-2.raw #36652 RT: 45.4207 min
FTMS, 760.3293@hcd30.00, z=+2, Mono m/z=760.32935 Da, MH+=1519.65141 Da, Match Tol.=0.02 Da

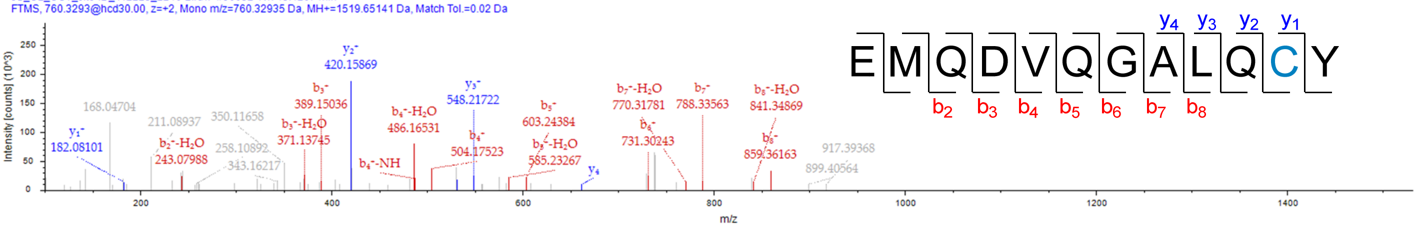


OGT1(517–524)

| #1 | b⁺ | b²⁺ | Seq. | y⁺ | y²⁺ | #2 |
|---|---|---|---|---|---|---|
| 1 | 138.06619 | 69.53673 | H | | | 8 |
| 2 | 195.08765 | 98.04746 | G | 897.44985 | 449.22856 | 7 |
| 3 | 309.13058 | 155.06893 | N | 840.42839 | 420.71783 | 6 |
| 4 | 422.21464 | 211.61096 | L | 726.38546 | 363.69637 | 5 |
| 5 | 660.29223 | 330.64975 | C-SAS_me... | 613.30140 | 307.15434 | 4 |
| 6 | 773.37629 | 387.19178 | L | 375.22381 | 188.11554 | 3 |
| 7 | 888.40323 | 444.70526 | D | 262.13975 | 131.57351 | 2 |
| 8 | | | K | 147.11280 | 74.06004 | 1 |

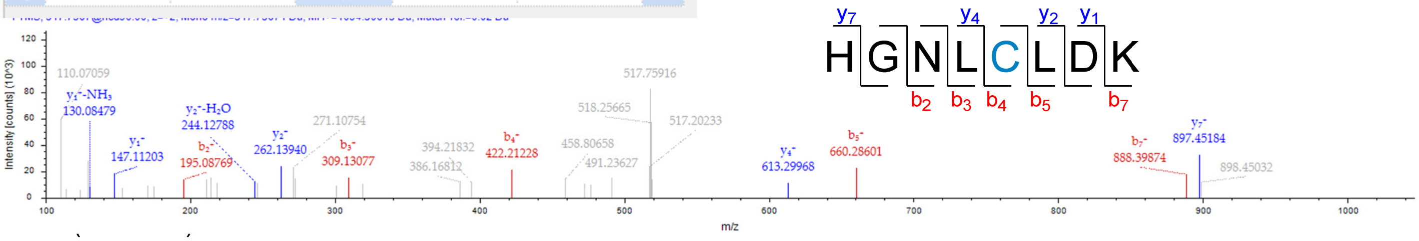


| #1 | b⁺ | b²⁺ | Seq. | y⁺ | y²⁺ | #2 |
|---|---|---|---|---|---|---|
| 1 | 157.10839 | 79.05783 | R | | | 13 |
| 2 | 214.12985 | 107.56856 | G | 1379.68751 | 690.34740 | 12 |
| 3 | 342.18843 | 171.59785 | Q | 1322.66605 | 661.83666 | 11 |
| 4 | 455.27249 | 228.13988 | L | 1194.60747 | 597.80737 | 10 |
| 5 | 526.30961 | 263.65844 | A | 1081.52341 | 541.26534 | 9 |
| 6 | 641.33655 | 321.17191 | D | 1010.48630 | 505.74679 | 8 |
| 7 | 740.40496 | 370.70612 | V | 895.45935 | 448.23331 | 7 |
| 8 | 978.48255 | 489.74491 | C-SAS_me... | 796.39094 | 398.69911 | 6 |
| 9 | 1091.56661 | 546.28694 | L | 558.31335 | 279.66032 | 5 |
| 10 | 1206.59355 | 603.80042 | D | 445.22929 | 223.11828 | 4 |
| 11 | 1307.64123 | 654.32425 | T | 330.20235 | 165.60481 | 3 |
| 12 | 1404.69400 | 702.85064 | P | 229.15467 | 115.08097 | 2 |
| 13 | | | L | 132.10191 | 66.55459 | 1 |

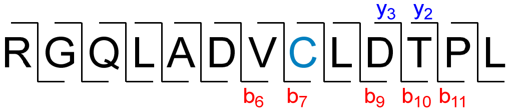


JS_CL_OGT_site-ID_112225_2240-2.raw #44797 RT: 52.5104 min
FTMS, 768.3964@hcd30.00, z=+2, Mono m/z=768.39642 Da, MH+=1535.78557 Da, Match Tol.=0.02 Da

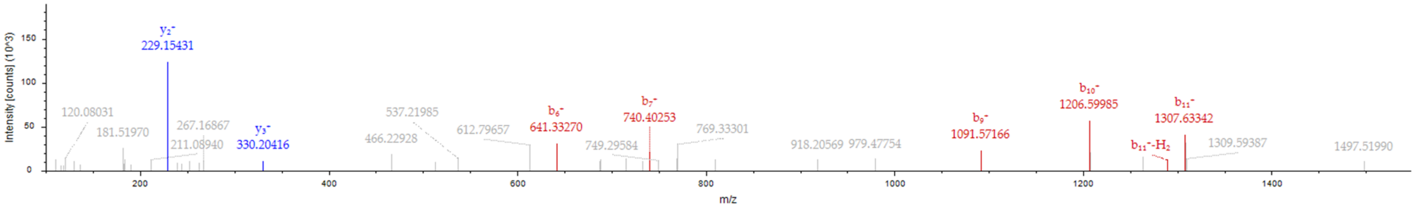


**Figure MS3.** Annotated MS/MS spectrum of labeled peptides with alkyne probe **E3.13** on OGT, including peptide coverage and labeled Cys407, Cys521, and Cys911.

AKT1(290–297)

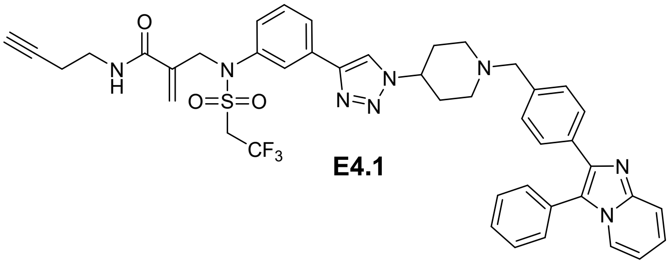

**E4.1**

| #1 | b⁺ | b²⁺ | Seq. | y⁺ | y²⁺ | #2 |
|---|---|---|---|---|---|---|
| 1 | 114.09134 | 57.54931 | I | | | 8 |
| 2 | 215.13902 | 108.07315 | T | 918.43895 | 459.72311 | 7 |
| 3 | 330.16596 | 165.58662 | D | 817.39127 | 409.19927 | 6 |
| 4 | 477.23438 | 239.12083 | F | 702.36433 | 351.68580 | 5 |
| 5 | 534.25584 | 267.63156 | G | 555.29592 | 278.15160 | 4 |
| 6 | 647.33990 | 324.17359 | L | 498.27445 | 249.64086 | 3 |
| 7 | 885.41749 | 443.21238 | C-SAS_me... | 385.19039 | 193.09883 | 2 |
| 8 | | | K | 147.11280 | 74.06004 | 1 |

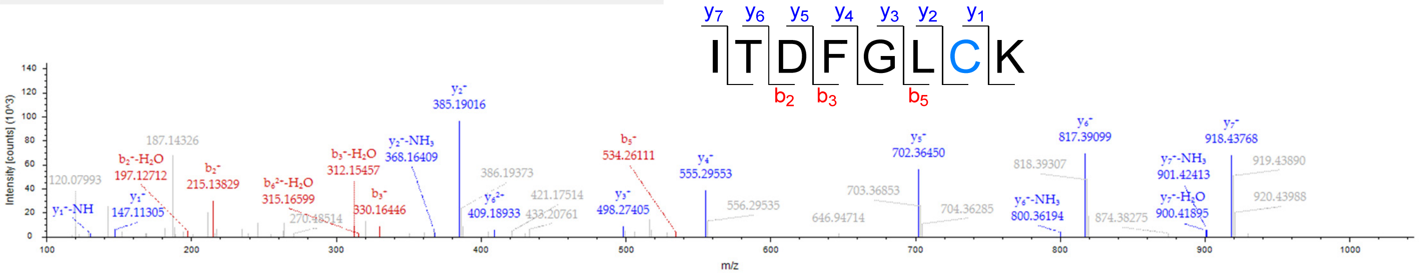


AKT1(308–328)

| #1 | b⁺ | b²⁺ | Seq. | y⁺ | y²⁺ | #2 |
|---|---|---|---|---|---|---|
| 1 | 102.05496 | 51.53112 | T | | | 21 |
| 2 | 249.12337 | 125.06532 | F | 2423.08599 | 1212.04663 | 20 |
| 3 | 487.20095 | 244.10411 | C-SAS_me... | 2276.01757 | 1138.51243 | 19 |
| 4 | 544.22242 | 272.61485 | G | 2037.93999 | 1019.47363 | 18 |
| 5 | 645.27010 | 323.13869 | T | 1980.91853 | 990.96290 | 17 |
| 6 | 742.32286 | 371.66507 | P | 1879.87085 | 940.43906 | 16 |
| 7 | 871.36545 | 436.18636 | E | 1782.81808 | 891.91268 | 15 |
| 8 | 1034.42878 | 517.71803 | Y | 1653.77549 | 827.39138 | 14 |
| 9 | 1147.51284 | 574.26006 | L | 1490.71216 | 745.85972 | 13 |
| 10 | 1218.54996 | 609.77862 | A | 1377.62810 | 689.31769 | 12 |
| 11 | 1315.60272 | 658.30500 | P | 1306.59099 | 653.79913 | 11 |
| 12 | 1444.64532 | 722.82630 | E | 1209.53822 | 605.27275 | 10 |
| 13 | 1543.71373 | 772.36050 | V | 1080.49563 | 540.75145 | 9 |
| 14 | 1656.79779 | 828.90253 | L | 981.42721 | 491.21725 | 8 |
| 15 | 1785.84039 | 893.42383 | E | 868.34315 | 434.67521 | 7 |
| 16 | 1900.86733 | 950.93730 | D | 739.30056 | 370.15392 | 6 |
| 17 | 2014.91026 | 1007.95877 | N | 624.27361 | 312.64045 | 5 |
| 18 | 2129.93720 | 1065.47224 | D | 510.23069 | 255.61898 | 4 |
| 19 | 2293.00053 | 1147.00390 | Y | 395.20374 | 198.10551 | 3 |
| 20 | 2350.02199 | 1175.51463 | G | 232.14042 | 116.57385 | 2 |
| 21 | | | R | 175.11895 | 88.06311 | 1 |

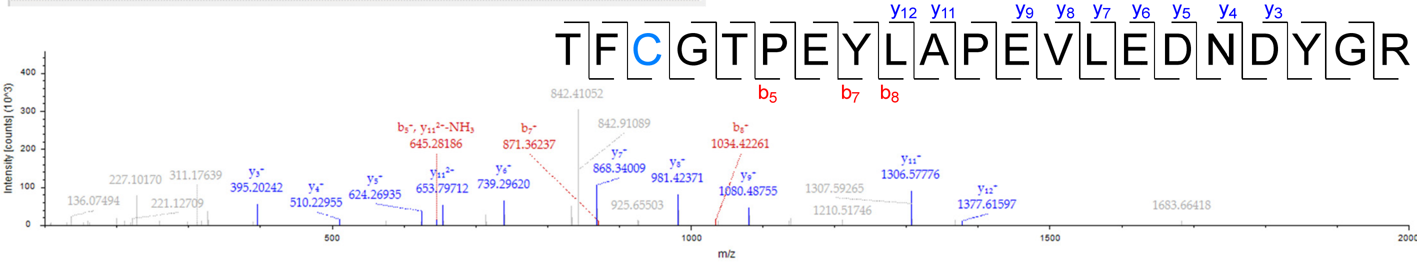


**Figure MS4.** Annotated MS/MS spectrum of labeled peptides with alkyne probe **E4.1** on AKT1**,** including peptide coverage and labeled Cys296 and Cys310.

Value Type: Theo. Mass [Da]

Ion Series | Neutral Losses | Precursor Ions | Internal Fragments

| #1 | b⁺ | b²⁺ | Seq. | y⁺ | y²⁺ | #2 |
|---|---|---|---|---|---|---|
| 1 | 129.06585 | 65.03657 | Q | | | 14 |
| 2 | 351.20268 | 176.10498 | K-SAS_NA... | 1490.78895 | 745.89811 | 13 |
| 3 | 450.27109 | 225.63918 | V | 1268.65213 | 634.82970 | 12 |
| 4 | 563.35515 | 282.18122 | I | 1169.58372 | 585.29550 | 11 |
| 5 | 620.37662 | 310.69195 | G | 1056.49965 | 528.75346 | 10 |
| 6 | 691.41373 | 346.21050 | A | 999.47819 | 500.24273 | 9 |
| 7 | 748.43520 | 374.72124 | G | 928.44107 | 464.72418 | 8 |
| 8 | 877.47779 | 439.24253 | E | 871.41961 | 436.21344 | 7 |
| 9 | 1024.54620 | 512.77674 | F | 742.37702 | 371.69215 | 6 |
| 10 | 1081.56767 | 541.28747 | G | 595.30860 | 298.15794 | 5 |
| 11 | 1210.61026 | 605.80877 | E | 538.28714 | 269.64721 | 4 |
| 12 | 1309.67867 | 655.34298 | V | 409.24455 | 205.12591 | 3 |
| 13 | 1472.74200 | 736.87464 | Y | 310.17613 | 155.59170 | 2 |
| 14 | | | K | 147.11280 | 74.06004 | 1 |

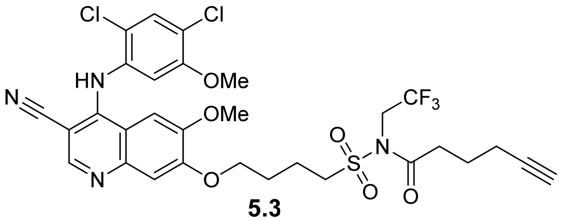

**5.3**

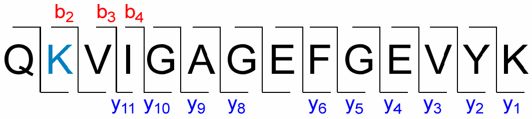


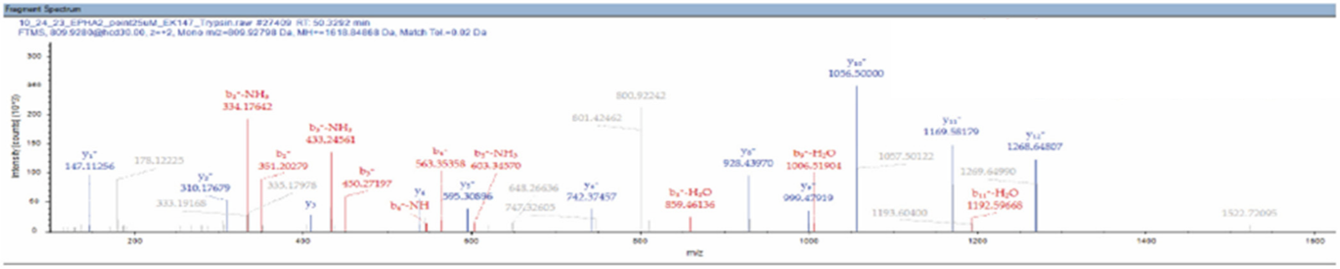


**Figure MS5.** Annotated MS/MS spectrum of a labeled peptide with alkyne probe **5.3** on EphA2**,** including peptide coverage and labeled lysine.

| #1 | b⁺ | b²⁺ | Seq. | y⁺ | y²⁺ | #2 |
|---|---|---|---|---|---|---|
| 1 | 116.03422 | 58.52075 | D | | | 17 |
| 2 | 229.11828 | 115.06278 | L | 1831.03643 | 916.02185 | 16 |
| 3 | 330.16596 | 165.58662 | T | 1717.95237 | 859.47982 | 15 |
| 4 | 477.23438 | 239.12083 | F | 1616.90469 | 808.95598 | 14 |
| 5 | 590.31844 | 295.66286 | L | 1469.83628 | 735.42178 | 13 |
| 6 | 812.45530 | 406.73129 | K-SAS_EK... | 1356.75221 | 678.87974 | 12 |
| 7 | 941.49790 | 471.25259 | E | 1134.61535 | 567.81131 | 11 |
| 8 | 1054.58196 | 527.79462 | L | 1005.57276 | 503.29002 | 10 |
| 9 | 1111.60342 | 556.30535 | G | 892.48869 | 446.74798 | 9 |
| 10 | 1212.65110 | 606.82919 | T | 835.46723 | 418.23725 | 8 |
| 11 | 1269.67257 | 635.33992 | G | 734.41955 | 367.71341 | 7 |
| 12 | 1397.73114 | 699.36921 | Q | 677.39809 | 339.20268 | 6 |
| 13 | 1544.79956 | 772.90342 | F | 549.33951 | 275.17339 | 5 |
| 14 | 1601.82102 | 801.41415 | G | 402.27110 | 201.63919 | 4 |
| 15 | 1700.88943 | 850.94836 | V | 345.24963 | 173.12845 | 3 |
| 16 | 1799.95785 | 900.48256 | V | 246.18122 | 123.59425 | 2 |
| 17 | | | K | 147.11280 | 74.06004 | 1 |

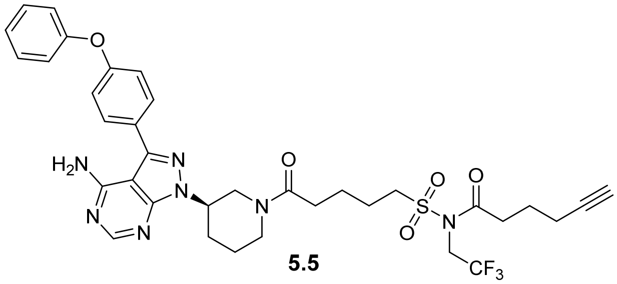

**5.5**

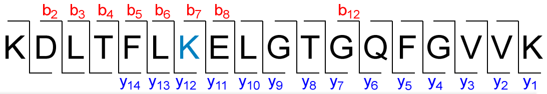


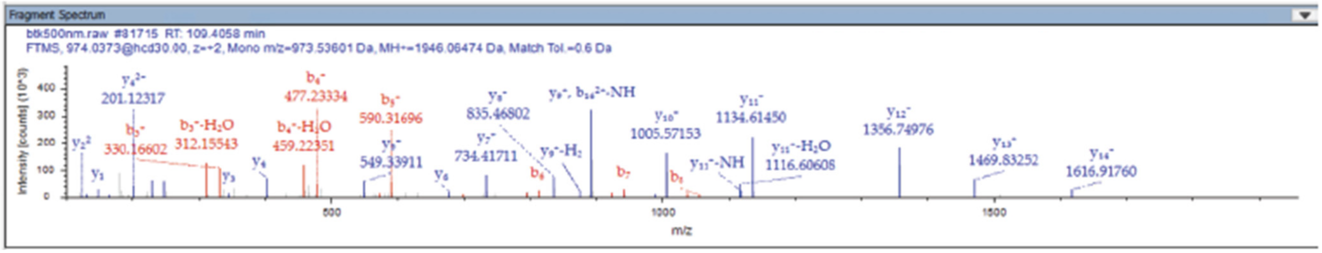


**Figure MS6.** Annotated MS/MS spectrum of a labeled peptide with alkyne probe **5.5** on BTK**,** including peptide coverage and labeled lysine.
